# The wiring logic of an identified serotonergic neuron that spans sensory networks

**DOI:** 10.1101/2020.02.24.963660

**Authors:** Kaylynn E. Coates, Steven A. Calle-Schuler, Levi M. Helmick, Victoria L. Knotts, Brennah N. Martik, Farzaan Salman, Lauren T. Warner, Sophia V. Valla, Davi D. Bock, Andrew M. Dacks

## Abstract

Serotonergic neurons modulate diverse physiological and behavioral processes in a context-dependent manner, based on their complex connectivity. However, their connectivity has not been comprehensively explored at a single-cell resolution. Using a whole-brain EM dataset we determined the wiring logic of a broadly projecting serotonergic neuron (the “CSDn”) in *Drosophila*. Within the antennal lobe (AL; first-order olfactory region), the CSDn receives glomerulus-specific input and preferentially targets distinct local interneuron subtypes. Furthermore, the wiring logic of the CSDn differs between olfactory regions. The CSDn innervates the AL and lateral horn (LH), yet does not maintain the same synaptic relationship with individual projection neurons that also span both regions. Consistent with this, the CSDn has more distributed connectivity in the LH relative to the AL, preferentially synapsing with principal neuron types based on presumptive transmitter content. Lastly, we identify protocerebral neurons that provide abundant synaptic input to the CSDn. Our study demonstrates how an individual modulatory neuron can interact with local networks and integrate input from non-olfactory sources.

## Introduction

Every neural network receives modulatory input from a variety of sources [1, 2], and in some cases from heterogeneous populations of neurons that release the same modulatory transmitter [3–5]. In mammals, one ubiquitous neuromodulator, serotonin, is released by tens of thousands to hundreds of thousands of neurons which originate in the raphe nuclei and project throughout the brain [6, 7]. Serotonergic raphe neurons are highly diverse in their projections, connectivity, and electrophysiological properties, and are implicated in a wide breadth of behaviors and physiological processes [4, 8–16]. Further, the raphe system receives monosynaptic input from up to 80 anatomical areas [8, 9]. As a result, a significant amount of work has focused on disentangling the functional and behavioral roles of serotonergic neurons. Several recent studies have suggested that serotonergic raphe neurons may be organized into functional subpopulations based on neuroanatomy, electrophysiology, and behavior [4, 12, 17–21]. For example, two parallel sub-systems of serotonergic raphe neurons collateralize complimentarily and are both activated by reward, yet have opposing responses to aversive stimuli and promote distinct behaviors [18]. Sparse neuron reconstructions in mice show that a single serotonin neuron can interconnect the olfactory bulb, piriform cortex, and anterior olfactory nucleus [19], demonstrating that a single serotonergic neuron can arborize several processing stages within the same sensory modality. Thus, determining the precise patterns of connectivity of single serotonergic neurons within and across the brain regions will be critical for understanding the seemingly heterogeneous effects of serotonergic systems.

Invertebrates are excellent models for studying the properties of individual neurons due to the wealth of identified neurons that can be consistently studied from animal to animal [22]. The fly olfactory system provides a numerically reduced model in which to study the detailed connectivity of an individual serotonergic neuron while minimizing inter-animal variability [23, 24]. Further, the genetic tractability and numerical simplicity of the fly enables the olfactory system to be studied at a single-cell resolution. Recently, a whole female adult fly brain electron microscopy volume (FAFB) was generated in which the connectivity of a neuron with its synaptic partners can be comprehensively determined at a single synapse resolution [25]. Taken together, these advantages position the fly as an ideal model to explore the nature of a single modulatory neuron’s synaptic connectivity between individual neurons and cell classes, as well as within and across brain regions.

Flies detect volatile chemicals via an array of olfactory receptor neurons (ORNs) housed in sensillum on their antennae. Each ORN expresses 1-2 differentially tuned chemosensory receptor proteins, and ORNs expressing the same receptor proteins converge within sub-compartments of the antennal lobe (AL; first-order olfactory processing neuropil) called glomeruli. Olfactory information is carried to second-order processing centers including the lateral horn (LH) and mushroom bodies (MB) by uniglomerular and multiglomerular projection neurons (uPNs and mPNs, respectively) along the medial AL tract (mALT) [26–29]. Synaptic communication between all principal olfactory neuron types in the AL is refined by diverse populations of local interneurons (LNs) that support an equally diverse set of network computations [30–39].

The AL and LH also receive input from a variety of extrinsic neurons, including two broadly projecting serotonergic neurons, the “contralaterally-projecting, serotonin-immunoreactive deutocerebral neurons” (CSDns) [23, 24]. The CSDns are the sole source of synaptic serotonin in the AL and LH, and receive synaptic input in both neuropils [40–43]. However, the CSDns are by no means uniform. CSDn active zone density varies across glomeruli [40] and a given odor can cause local excitation of CSDn branches in the LH, yet widespread inhibition in the AL [44]. This suggests that local synaptic connectivity supports different coding schemes across olfactory processing neuropil for this single serotonergic neuron [42, 44]. Furthermore, different subsets of principal olfactory neurons express distinct 5-HT receptor subtypes [45]. Finally, the behavioral function of the CSDns varies with the odor identity and concentration tested, in some cases implying a suppression of odor-guided behavior and in others an enhancement [41, 46, 47]. Taken together, these studies suggest that even a single modulatory neuron can have heterogeneous connectivity within the networks that it targets. However, the organizing principles of the connectivity of single serotonergic neurons have not been comprehensively determined. Specifically, it is unclear how the connectivity of a single serotonergic neuron varies between specific neuron classes, within a neural network, or even across the different networks that it innervates.

To comprehensively study the connectivity of the CSDn at a single cell resolution, we used the FAFB electron microscopy volume [25] to fully reconstruct the CSDn in the AL and LH, identified individual synaptic partners across neuropil, and generated comprehensive connectomes of the CSDn within select glomeruli. While some coarse anatomical features of the CSDn are consistent across neuropil, we find that the overall connectivity of the CSDn is highly complex, as it differs within the AL as well as between the AL and LH. We show that within the AL, the CSDn targets local processing networks: it synapses extensively and reciprocally with subsets of local interneurons and receives glomerulus-specific input from principal olfactory neuron types. This pattern of organization is not conserved in the LH, where the connectivity of CSDn with principal LH cell types varies with transmitter content of synaptic partners. Lastly, we demonstrate that the CSDn receives previously undescribed top-down input from three populations of extrinsic neurons, including a population of mechanosensory neurons. Taken together, we establish a wiring logic in which a single serotonergic neuron selectively targets specific cell types within a sensory network, while integrating heterogeneous, reciprocal input from local networks and top-down input from extrinsic sources. This heterogeneity at the level of a single cell may therefore contribute an additional degree of complexity to the role of populations of modulatory neurons.

## Results

### General morphological features of the CSDn are conserved across brain regions

Our first goal was to systematically compare broad morphological characteristics such as branching density and distribution of synaptic sites of a single CSDn across and within brain regions. The somata of the two CSDns reside in the lateral cell cluster of each AL (Figure 1A). A single CSDn produces sparse processes on its primary neurite within the ipsilateral AL, then follows the medial AL tract (mALT) to the ipsilateral protocerebrum where it innervates the antler, superior lateral protocerebrum (SLP), mushroom body calyx (MBC), and lateral horn (LH). The CSDn then crosses the midline to innervate these same protocerebral regions on the contralateral side of the brain. Finally, the CSDn projects along the contralateral mALT to densely innervate the contralateral AL (Figure 1A, B).

**Figure 1:**
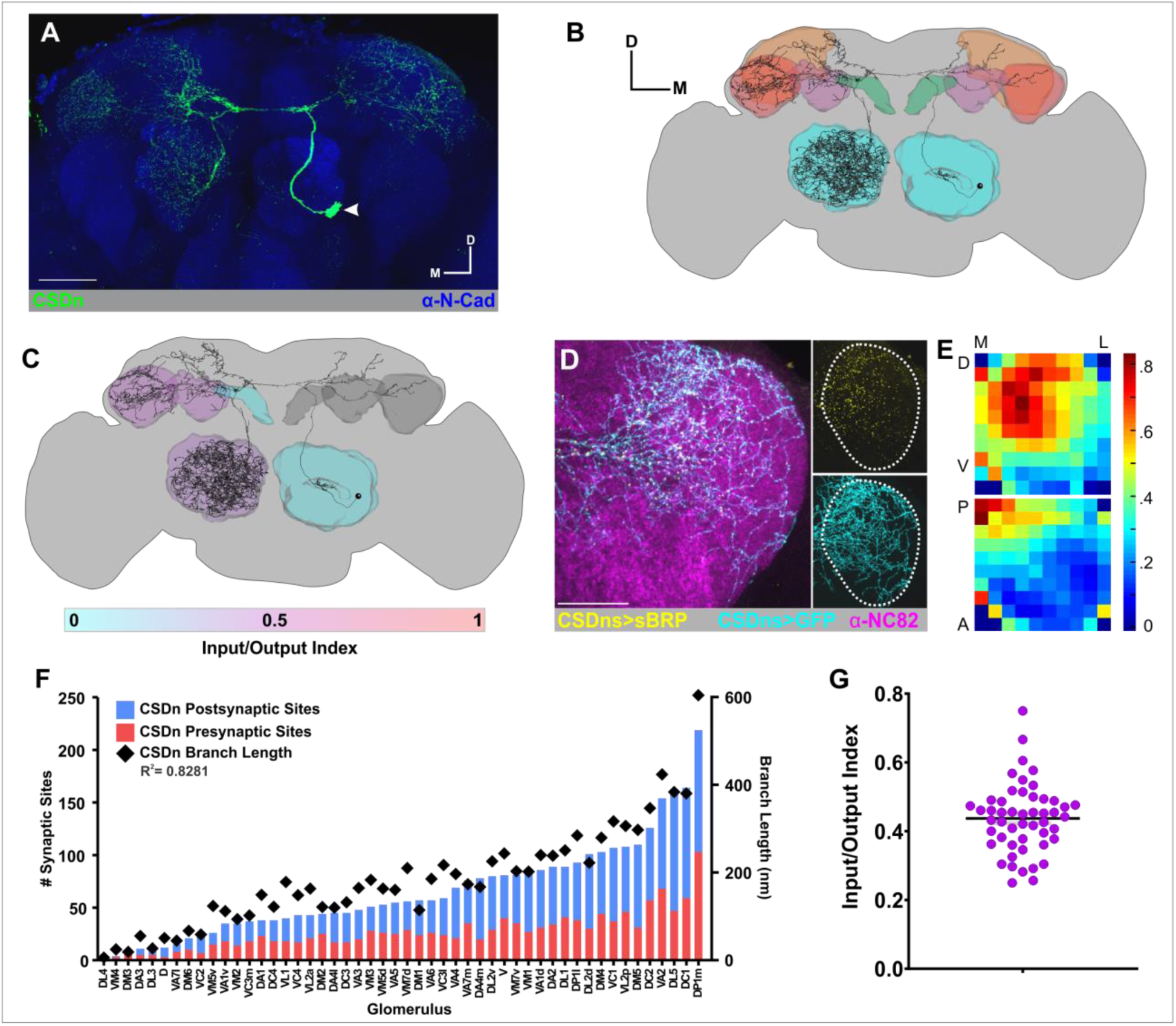
CSD_n_ EM reconstruction and distribution of synaptic sites. (A) Multi-color flip out (green) highlights the arborization pattern of a single CSDn. N-Cadherin labeling delineates neuropil (blue). (B) Reconstruction of the left-hand CSDn (soma in the fly’s left hemisphere) from the FAFB dataset. (C) The CSDn has a mix of input and output sites across most olfactory neuropil. Shading based on the “lnput/Output Index” of the CSDn was calculated as the # inputs/(# inputs + # outputs) where 0 = output only and 1 = input only. (D) Expression of Brp-short puncta (yellow) labels CSDn active zones in the lateral horn. GFP (cyan) labels the CSDn’s arborizations and NC82 delineates neuropil (magenta). Scale bar = 50 um. (E) Heatmaps of the distribution of Brp-short puncta (i.e. CSDn active zones) in the LH. Puncta density is higher in the mediodorsal and posterior regions of lateral horn (top and bottom heat maps, respectively). (F) Glomeruli rank-ordered by the total number of CSDn synaptic sites. The number of CSDn presynaptic sites (red bars) and postsynaptic sites (blue bars) are correlated to total CSDn cable length in each glomerulus (black diamonds), R^2^=0.8231. (G) The Input/Output Index of the CSDn in each glomerulus is fairly consistent across glomeruli. Mean = 0.4359, SEM = 0.01, Coefficient of Variation = 22.96%.

Due to the intricacy and size of the CSDn, we focused our efforts on reconstructing a single CSDn in the right hemisphere of the brain (i.e. CSDn soma in the fly’s left hemisphere, the “left-hand CSDn”) where most neuronal reconstruction in the FAFB dataset [25, 48–50] has occurred. We manually reconstructed over 23 x 10^6^ nm of cable which includes a complete reconstruction of the CSDn and its synaptic sites in the ipsilateral AL, contralateral AL, contralateral MBC and contralateral LH (Figure 1B). We used NBLAST [51] to geometrically compare our tracing to a skeletonized CSDn from a light microscopy image dataset [52] and obtained a similarity score of 0.716, indicating a match (where 1 equals perfect alignment with dataset) (Figure 1 – figure supplement 1A). Despite functional differences in the organization of the AL and LH networks [53, 54], broad features of CSDn morphology and synapse distribution are similar across them. CSDn cable length per neuropil volume is similar in the AL (45.284 nm/um^3^) and LH (38.832 nm/um^3^) respectively, despite the CSDn innervating the AL much more extensively than the LH, with 17 x 10^6^ nm of cable compared to 6 x 10^6^ nm in the LH. We did find, however, that the CSDn is sparser in the MBC, with a total of 0.9 x 10^6^ nm of cable length and branching density of 8.792 nm/um^3^ (Figure 1 – figure supplement 1C).

Expanding upon previous studies using transgenic markers [24, 40–42], we found that the CSDn has input and output sites mixed along its neurites in all olfactory neuropil, except for the ipsilateral AL and protocerebral region called the “antler” (Figure 1B) which are postsynaptic. To determine if the ratio of pre- and post-synaptic sites of the CSDn differs across the contralateral AL, LH, and MBC, we calculated an “input/output index” (# presynaptic sites / (# presynaptic + # postsynaptic sites). All three neuropils have relatively similar input/output indices (ranging from 0.42-0.46; Figure 1C, Figure 1 – figure supplement 1C) and synaptic density (ranging from 0.76-1.02 synapses/10 um; Figure 1 – figure supplement 1C). Taken together, this indicates that coarse traits, like total cable length and input/output index, are consistent across each olfactory network.

Within individual brain regions, however, the CSDn has non-uniform projections [24, 40, 47] and distribution of presynaptic sites. The number of CSDn presynaptic sites varies between AL glomeruli, yet is consistent for a given glomerulus across animals [40]. Furthermore, the distribution of the active zone marker Brp_short_ [55–58] is most dense in the mediodorsal and posterior regions of the LH (Figure 1D, E). We therefore sought to determine if the balance of pre- and postsynaptic sites of the CSDn in the AL is also consistent across glomeruli and if the absolute number of pre- and postsynaptic sites scale with branch length as it does for global measures from the AL, LH, and MBC. We chose to focus on the AL because of its glomerular organization which allows us to easily compare discrete subregions as opposed to the LH and MBC in which subregions are less easily discernable. Within each glomerulus, we quantified the number of pre- and postsynaptic sites of the CSDn and calculated its branch length. We found that while the total number of pre- and postsynaptic sites on the CSDn varies across glomeruli, the “input/output index” for each glomerulus is fairly consistent (Figure 1F,G, Mean = 0.437, SEM = 0.014, Coefficient of Variation = 22.96%), and the number of pre- and postsynaptic sites scale with branch length (R^2^ = 0.924 and 0.861 for pre- and postsynaptic sites respectively; Figure 1 – figure supplement 2A,A’). We also found that the synaptic density across all glomeruli is significantly less variable for presynaptic sites than postsynaptic sites (Levene’s test for homogeneity of variance, p<0.005; Figure 1-figure supplement 2B). Finally, we found a weak correlation between branch length and glomerular volume (Figure 1 – figure supplement 2C), indicating that the density of CSDn pre- and postsynaptic sites is consistent, based on branch length and that glomerulus specific differences in synaptic density are due to density of innervation.

### CSDn connectomes across glomeruli are distinct

The variation in CSDn pre- and postsynaptic density suggests that the specific neurons with which the CSDn interacts may also vary across glomeruli. In the AL, the composition of the neuronal population within each glomerulus (the “demographics”) correlates with the odor tuning of the ORNs that innervate that glomerulus [59]. Glomeruli that receive input from more narrowly tuned ORNs are innervated by more PNs relative to LNs, and glomeruli that receive input from more broadly tuned ORNs are innervated by more LNs. Furthermore, the sensitivity of PNs to GABAergic inhibition from LNs is inversely correlated to odor tuning [31], consistent with narrowly glomeruli receiving less input from LNs. Since the CSDn receives input from ORNs and PNs in a subset of glomeruli [40], we sought to systematically explore the differences in glomerulus-specific demographics of its synaptic partners (Figure 2). We reconstructed all of the CSDn’s presynaptic and postsynaptic partners to identification within 9 glomeruli: DA1, DA2, DC1, DM1, DM5, VC2, VM1, VM2, and VM3 (Figure 2 – figure supplement 2,3). These glomeruli were chosen because they vary in their odor-tuning (as measured by lifetime sparseness; [59]), number of CSDn synaptic sites, and CSDn branch length (Figure 2 – figure supplement 1), thus are likely representative of AL glomeruli. Thus we could determine if there is an obvious logic behind the synaptic connectivity of the CSDn across glomeruli.

**Figure 2:**
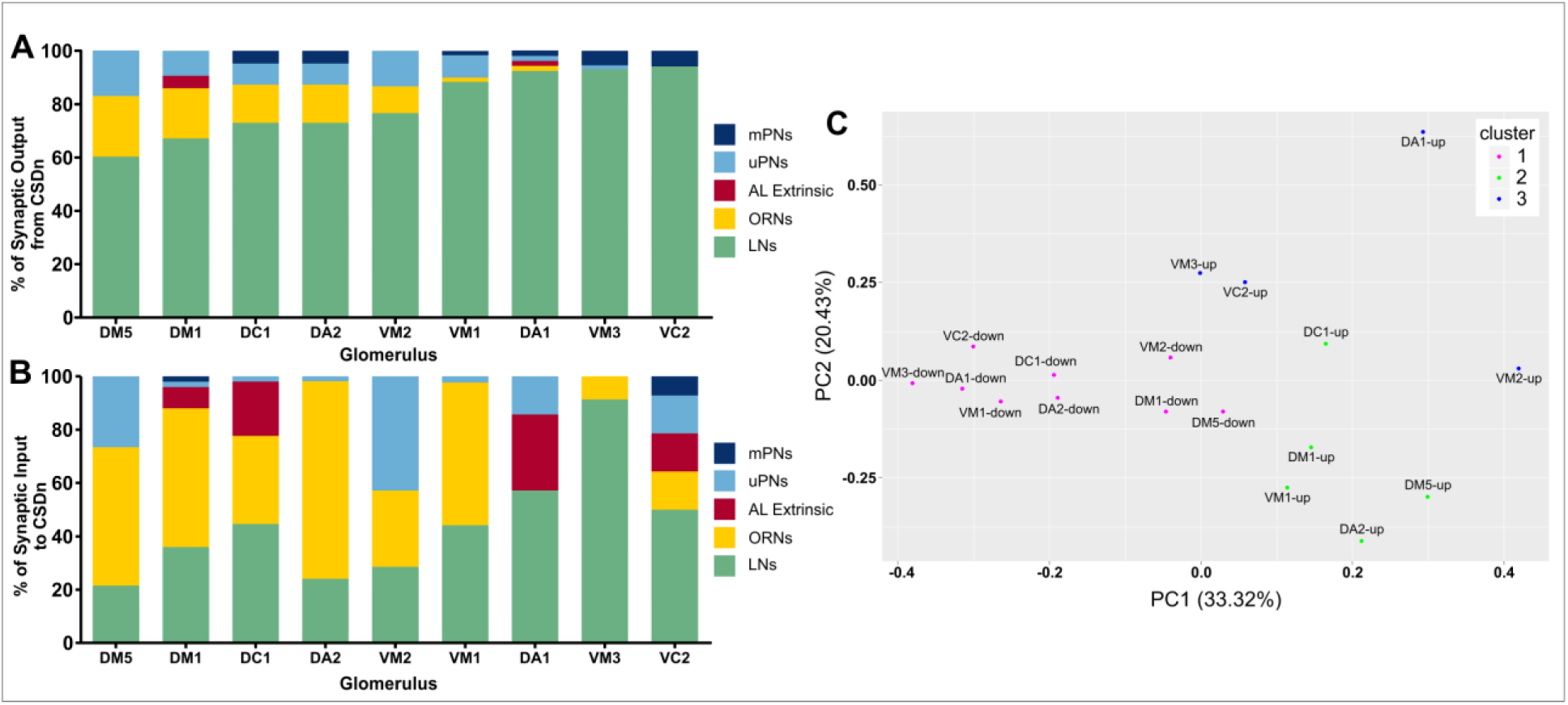
The CSDn has distinct connectivity across glomeruli. (A) Percent of input from the CSDn onto all postsynaptlc partners reconstructed in 9 glomeruli. The CSDn predominantly targets LNs across all glomeruli. Glomerulus order based on ranked ordered amount of Input from the CSDn to LNs. (B) Percent of Input to the CSDn from all presynaptic partners reconstructed in 9 glomeruli. Composition of presynaptic targets within each glomerulus is more diverse than for CSDn output. (C) Synapse fractions segregate into 3 clusters in PC space, based on whether the fraction is for upstream partners (magenta) or downstream partners (green and blue) based on k-means clustering.

Despite the large differences in glomerulus-specific innervation density of the CSDn (Figure 1F), the demographics of CSDn downstream synaptic partners are relatively uniform across glomeruli (Figure 2A). Although the percent of output from the CSDn onto ORNs, PNs, and extrinsic neurons in the 9 glomeruli varies, we found that 60%-94% of the CSDn’s downstream synaptic partners are LNs, regardless of odor tuning, and that lifetime sparseness does not correlate with CSDn output to LNs (R^2^=0.003, p=0.905; Figure 2 – figure supplement 4A). For example, over 90% of the postsynaptic partners of the CSDn in DA1 and VM3 are LNs. However, DA1 has a lifetime sparseness of 0.98 and VM3 has a lifetime sparseness of 0.56 indicating that the downstream targets do not vary with tuning breadth of a given glomerulus. However, the proportion of CSDn synapses onto ORNs varies inversely with the proportion of CSDn synapses upon LNs (R^2^=0.98, p<0.001; Figure 2 – figure supplement 4B).

In contrast, the demographics of upstream synaptic partners to the CSDn in each glomerulus are highly variable (Figure 2B). The neuron types which provide input to the CSDn do not correlate to glomerulus-specific CSDn innervation density or lifetime sparseness of the cognate ORNs. For example, VM2 and VM3 are both broadly tuned glomeruli (Figure 2 – figure supplement 1). However, LNs provide the largest fraction of input to the CSDn in VM2, whereas uPNs provide the most input in VM3. The CSDn receives most of its input from ORNs in both DA2 and VM1, which are narrowly tuned, yet in DA1 which is also narrowly tuned, the CSDn receives most of its input from LN subtypes (see below). We ran a principal component analysis (PCA) in which the first component is largely explained by opposing vectors of vLNs and ORNs and PC2 is largely explained by opposing vectors of “uncategorized LNs” and uPNs. However, we did not find that the demographics of synaptic partners covary as their eigenvectors are largely distributed throughout the PCA (Figure 2 – figure supplement 5A).

The high apparent degree of variability in the demography of input to the CSDn is consistent with our observation that the glomerulus-specific density of postsynaptic CSDn sites is more variable relative to the density of CSDn presynaptic sites (Figure 1 – figure supplement 2B). A PCA including all of the glomerulus specific sets of upstream and downstream partners (Figure 2C) confirms that the composition of upstream partners is more variable, as the mean distance between coordinates for upstream partner sets is significantly higher than between downstream partners sets (Figure 2 – figure supplement 5D; p=0.0007; Student’s t-test). In addition, the synapse fractions of downstream partner sets cluster together separately from the upstream partner sets which form two sub-clusters (Figure 2C; Figure 2 – figure supplement 5B). This indicates that the CSDn has distinct patterns of synaptic output and input across glomeruli. Overall, while the CSDn predominantly influences LNs, glomerulus specific input likely tempers this influence depending upon the odor that the fly encounters.

### The CSDn has distinct connectivity with LN types

We found that the CSDn predominantly targets LNs within the nine glomeruli in which we reconstructed the CSDn’s synaptic partners. However, LNs are extremely heterogeneous in their morphology, physiology, and transmitter profile [30-32, 34, 35, 37-39, 60-62]. Thus, to establish a systematic framework for the connectivity of the CSDn to LNs, we determined the LN subtypes to which the CSDn is preferentially connected. The CSDn synapses with at least 84 LNs throughout the AL. Previous work has categorized LNs based on their morphology and the number of glomeruli that they innervate into the following subtypes: panglomerular, all but a few (“ABAF”), continuous, patchy, oligoglomerular, LNs with somata in the subesophageal zone (SEZ) and ventral [30, 61, 63]; Figure 3A-E). Although we did not find evidence for the CSDn being strongly connected synaptically (>10 synapses) to oligoglomerular or panglomerular LNs, these LN types could be partially reconstructed neurons within the group of uncategorized LNs (see below) to which the CSDn is weakly connected but we did not classify further.

**Figure 3:**
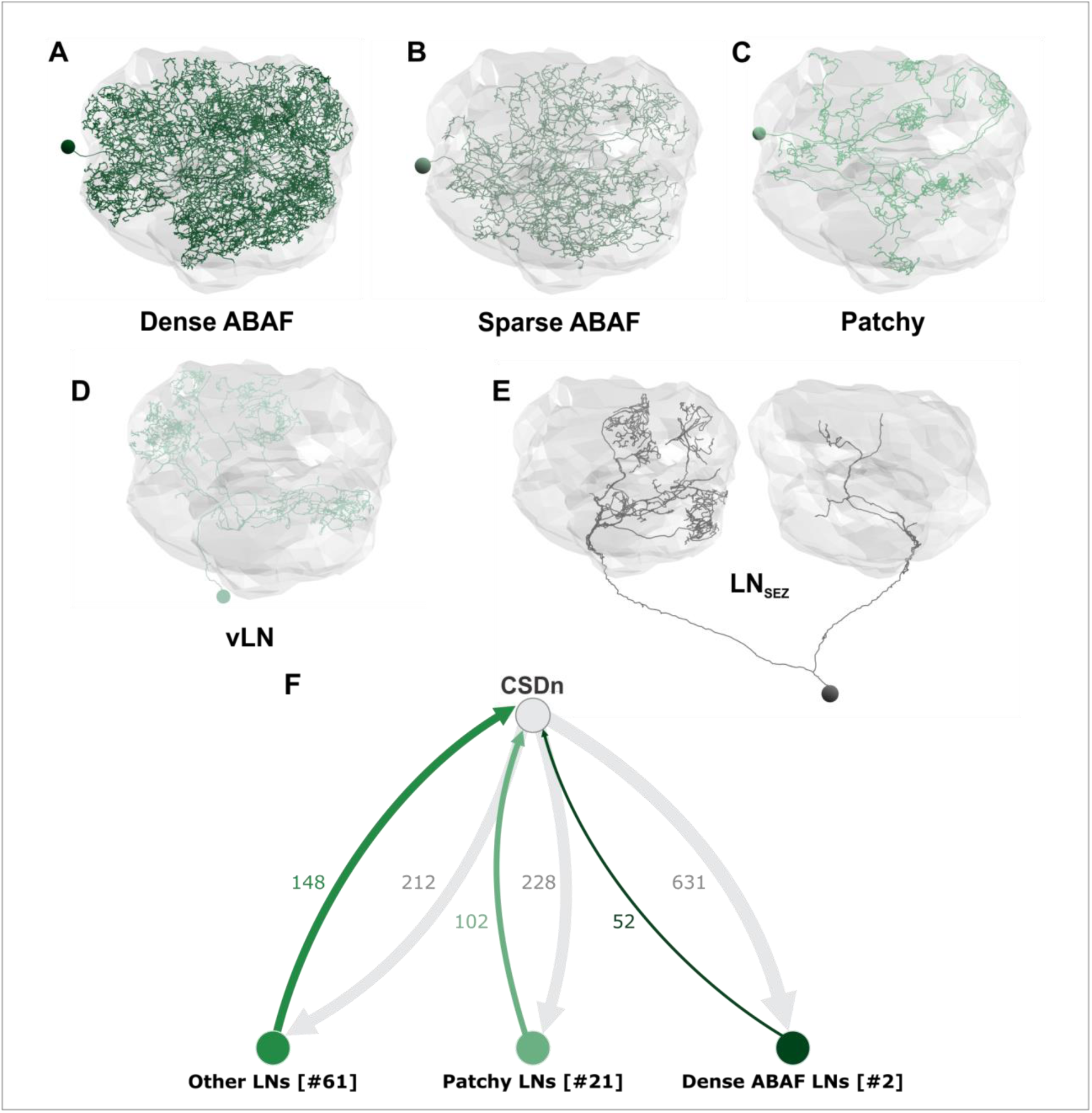
The CSDn has distinct connectivity with LN sub-populations. Examples of LNs with connectivity to the CSDn. (A) Dense ABAF LNs innervate ∼50 glomeruli. (B) The Sparse ABAF also innervates ∼50 glomeruli but has far less branchpoints compared to the dense ABAF. (C) Patchy LNs characteristic looping structure within the glomeruli that they innervate. (D) vLNs have their soma ventral to the AL and are likely glutamatergic. (E) AL projecting SEZ LNs”(LN_SEZ_) have their somata ventral and project bilaterally to both ALs. These LNs resemble the Keystone LNs reconstructed in a larval EM dataset (Berck et al., 2016). (F) The CSDn is most strongly connected to two Dense ABAF LNs as well as Patchy LNs. The “Other LNs” group include LN_SEZ_s, vLNs, and otherwise uncategorized LNs. “[#]” indicates number of neurons within a population. Synaptic connections derive from the 9 completely reconstructed glomeruli in Figure 2 and synaptic connections from other glomeruli found in the course of reconstructing the LN.

Despite the diversity of LN types, we found that 50-80% of the CSDn’s output to LNs is directed onto two subtypes of LNs across the 9 glomeruli; two densely branching ABAF LNs which each innervate 48 and 49 glomeruli, respectively (Figure 3 – figure supplement 2) and patchy LNs which each innervate ∼30 glomeruli [30]. The CSDn provides over 600 synapses to the two dense ABAF LNs, approximately one-third of its synaptic output within the AL and receives input from the ABAFs at over 50 synaptic sites (Figure 3F; Figure 3 - figure supplement 2). We identified and reconstructed another ABAF LN which innervates at least 44 glomeruli but branches far less extensively than the two densely branching ABAF LNs. Using a K-nearest neighbor analysis (KNN) we calculated the distribution of distances between branch points of a neuron and its neighboring branch points (see Methods). We found that the branch point distributions of the two dense ABAF LNs closely overlap, in contrast to the sparse ABAF LN which has a much larger range of branch point distributions and thus has far less overlap with the dense ABAF (Figure 3 – figure supplement 3). This indicates that the dense and sparse ABAF LNs indeed represent two separate sub-classes. Moreover, this sparse ABAF LN has little connectivity to the CSDn (Figure 3 - supplement 2A).

The CSDn is also highly connected to the patchy LNs which innervate sub-volumes, or “patches”, of individual glomeruli, and as a population tile across the entirety of the AL [30]. Before innervating a given glomerulus, patchy LNs have long, highly looping processes that lack pre- or postsynaptic sites, which further assisted in their identification. Although the biological significance of these convoluted processes is unclear, similar examples have been reported in *Drosophila* and other invertebrates [64–66]. Once within a glomerulus, a patchy LN branches extensively within a subregion of a glomerulus and produces many pre- and postsynaptic sites (Figure 3C; [30]). In contrast to the dense ABAF, the CSDn has 2:1 reciprocal connectivity to a population of at least 21 patchy LNs (Figure 3F). KNN analysis supports the hypothesis that these patchy LNs belong to the same morphological subclass (Figure 3 – figure supplement 3C). For the remaining LNs, the CSDn has weak (<5 synapses) reciprocal synaptic connectivity with ventral LNs (Figure 3D; Figure 3 – figure supplement 1C), which are most likely glutamatergic [32, 61], as well as a pair of bilaterally projecting LNs whose somas are in the SEZ (“AL projecting SEZ LNs”; LN_SEZ_), similar to the “Keystone” LNs (Figure 3E) upon which the CSDn synapse in the larval *Drosophila* AL [63]. An additional 46 unclassified LNs which were reconstructed from the 9 glomeruli are grouped as “uncategorized LNs” as they are not classified as ABAF, LN_SEZ_s, patchy, or vLNs (Figure 3F; Figure 3 – figure supplement 1C). Collectively, the CSDn provides at least 212 synapses to and receives at least 148 synapses from these LNs that are not ABAFs or patchy LNs, although they are likely not a monolithic group. Taken together, these results suggest that while LNs are the major target of the CSDn in the AL, the CSDn preferentially targets one particular LN subclass, the dense ABAF LNs, and has reciprocal connectivity with the larger population of patchy LNs (Figure 3 – figure supplement 1B).

### CSDn connectivity varies across olfactory processing regions

Individual serotonin neurons in mice and flies alike span several stages of olfactory processing [19, 23, 40], however it is unclear if the wiring logic of individual serotonergic neurons is conserved across brain regions. For instance, do serotonergic neurons interact with the same individual neuron across multiple brain regions? Are general connectivity rules maintained from one processing stage to the next? We began by asking if the connectivity of the CSDn to individual PNs differs across the AL and LH as both the CSDn and PNs span both brain regions. Previously published PN reconstructions [25, 50, 67] were grouped into three subtypes based on whether they were uniglomerular (uPN) or multiglomerular (mPN), and the AL tract along which the mPNs project, either mALT or mlALT [50]. In addition to the AL tract along which they project, mPNs functionally differ from each other in terms of the transmitter they express with mALT mPNs being cholinergic and mlALT mPNs being GABAergic [50, 68–70]. Collectively, 78 mALT uPNs (of 149), 46 mALT mPNs, and 28 mlALT mPNs (of 197 total mPNs) are synaptically connected to the CSDn across the AL and LH (Figure 4A-C). This is likely an underestimation of the total CSDn:PN connectivity in the AL, as not all PN dendrites were reconstructed to completion. However, any PN branches that synapse with the CSDn in the 9 glomeruli that we sampled were reconstructed to the primary process of the neuron (i.e. to identification).

**Figure 4:**
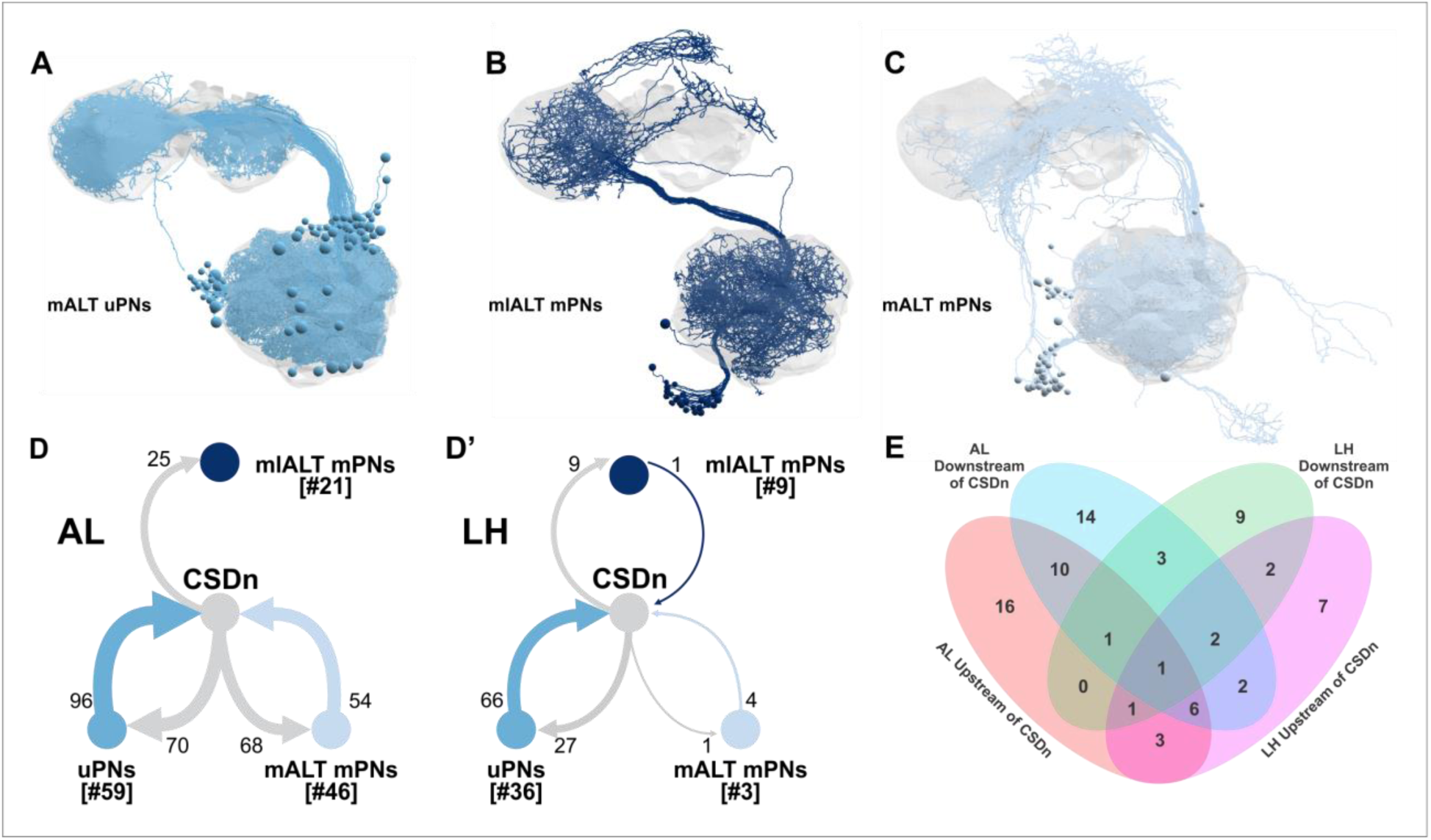
CSDn Connectivity with PNs in the AL and LH. EM reconstructions of PNs with connectivity to the CSDn (A) mALT uPNs, (B) mlALT mPNs, and (C) mALT mPNs. Number of synaptic connections of PN subtypes with the CSDn In the AL (D) and LH (D’). (E) uPNs have varied connectivity with the CSDn across the AL and LH. Number represents number of neurons, rather than synapse counts.

We found that the three PN types have distinct connectivity with the CSDn from one another both within and across the AL and LH. For instance, of the different PN subtypes, the CSDn is most strongly connected to the uPNs as a population, maintaining reciprocal connectivity with 3:2 uPN:CSDn ratio in both the AL and LH (Figure 4D, D’). While most individual uPNs are weakly connected to the CSDn (<10 synapses), a few individual uPNs are strongly connected. In particular, two DM5 uPNs have 16-18 synapses onto the CSDn in the AL, consistent with previous work using transgenic and physiological approaches demonstrating that DM5 PNs synapse with the CSDn [40]. The mALT mPNs also have balanced reciprocal synaptic connectivity with the CSDn in the AL, however, they have almost no connectivity with the CSDn in the LH (Figure 4D, D’). Finally, the CSDn has unidirectional connectivity with the mlALT mPNs in the AL with the CSDn providing input to, but not receiving input from them (Figure 4D; Figure 4 – figure supplement 2). Thus, similar to LNs, the connectivity of the CSDn with PNs varies across PN subtypes and varies between the AL and the LH. Even within a given glomerulus individual uPNs have different connectivity with the CSDn. For example, of the five DA2 uPNs, one synapsed upon the CSDn in the AL, two received synaptic input from the CSDn in the AL, one is reciprocally connected to the CSDn in the LH, and a fifth receives input from the CSDn in both brain regions (Figure 4 - figure supplement 1). The uPNs from the same glomerulus can differ in their connectivity within the AL [66] and MBC [71], and it appears that this feature extends to the CSDn as well.

To determine if the CSDn maintains its connectivity to individual neurons across brain regions, we asked if the CSDn is synaptically connected to the same individual uPNs in both the AL and LH. Although the CSDn could be connected to a given PN within both the AL and the LH, the connectivity relationship of a given uPN is rarely conserved across these processing stages (Figure 4E). For instance, in the AL there are 21 uPNs that receive synaptic input from the CSDn but are not reciprocally connected. Of these 21 uPNs, only 2 maintain this relationship in the LH. Thus, the CSDn has heterogeneous synaptic connectivity to individual neurons across processing stages of the olfactory system. Furthermore, a single serotonergic neuron can be differentially connected to “equivalent” neurons within (i.e. uPNs from the same glomerulus) and between multiple processing stages.

Finally, we sought to determine if broad features of CSDn connectivity to principal neuron types in the AL are maintained for the principal neuron types of the LH. While the CSDn primarily targets LNs within the AL, it has extensive connectivity with a diverse set of neurons within the LH in addition to PNs. Although we did not comprehensively reconstruct all of the synaptic partners of the CSDns within the LH, we were able to leverage a rich dataset of previously traced neurons [48–50] to subsample the populations of cells to which the CSDns are connected. There are at least 82 cell types of lateral horn neurons [48] which can be classified into three broad anatomical types; LH output neurons (LHONs), LH local neurons (LHLNs) and LH input neurons (LHINs). These can be further subdivided based on cell body cluster location (anterior ventral; AV, anterior dorsal; AD, posterior ventral; PV and posterior dorsal; PD) and their expression of acetylcholine, GABA, or glutamate [48–50]. Both CSDns are synaptically connected to neurons from all three categories (Figure 5A-D) and the strength of specific relationships, as well as the degree to which each relationship is symmetrical, varies with putative transmitter (Figure 5 – figure supplement 1). The CSDns are most strongly reciprocally connected to cholinergic LHONs, GABAergic LHLNs, and glutamatergic LHLNs (Figure 5E, F). Although the CSDns have mostly sparse (between 1-10 synapses) connectivity to individual LH neurons, the CSDns are strongly connected to individual LHINs, receiving strong synaptic input from two cholinergic LHINs and providing strong synaptic input to a GABAergic LHIN (Figure 5C, E, F). Thus, the general connectivity logic of the CSDn appears to differ between the AL and LH. While the CSDn connectivity in the AL is highly biased towards LNs, it has more distributed connectivity across principal cell types in the LH, with some exceptions, such as individual LHINs.

**Figure 5:**
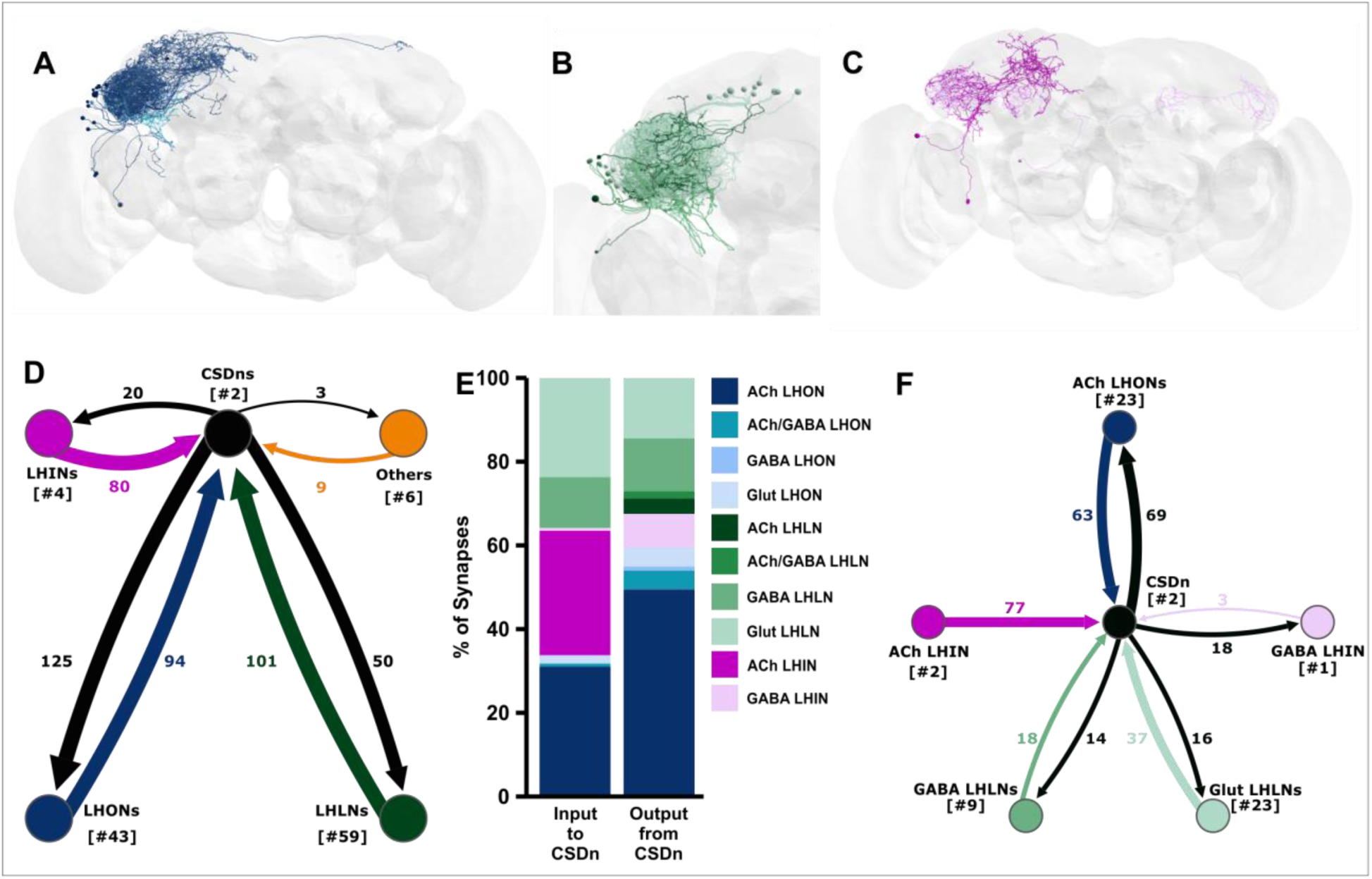
CSDn Connectivity with Lateral Horn Neurons. EM reconstruction of (A) LHONs, (B) LHLNs, and (C) LHINs that have connectivity with both CSDns. Colorization based on transmitter content (see E). (D) The CSDn reciprocally connectivity to each class of LH neurons. (E) Percent of input to/from the CSDns onto populations of lateral horn neurons. The CSDns have the most connectivity to and from cholinergic LHONs followed by glutamatergic and GABAerglc LHLNs. (F) Connectivity of LH neuron types that are strongly connected based ontransmltter content.

### The CSDn receives abundant synaptic input from distinct populations of protocerebral neurons

While CSDn processes have a mixture of pre- and postsynaptic sites in olfactory neuropils, CSDn processes are almost purely postsynaptic within a protocerebral region called the antler (ATL) (Figure 2C). We next asked what neurons provide strong synaptic input to the CSDn in the ATL. We reconstructed neurons that provide synaptic input to the CSDn in the ATL and identified a population of 10 morphologically similar neurons that collectively provide over 240 synapses to both CSDns in the ATL, LH, and several other protocerebral regions (Figure 6A; Figure 6 – figure supplement 1B). These neurons have their somata near the LH, project ventrally into the ipsilateral wedge where they branch extensively before projecting back dorsally, crossing the midline and then projecting ventrally into the contralateral wedge. The descending processes in each hemisphere project anteriorly into the anterior ventrolateral protocerebrum and medially into the saddle (Figure 6 – figure supplement 1C,D,E). These wedge projection neurons (WPNs) are morphologically similar to a population of unilaterally projecting WPNs [72] and have also been described in prior analyses of clonal units [73]. Due to their projections to both brain hemispheres, we refer to these WPNs as “Bilaterally projecting WPNs” (WPN_B_s). Consistent with a role in processing mechanosensory input to the antennae, the WPN_B_s receive a large amount of synaptic input within the lateral portion of the ipsilateral wedge (Figure 6-figure supplement 1C’,D’,E’), a second-order mechanosensory center [72, 74–78]. It should be noted that there are more than 10 WPN_B_s in total as we did identify other WPN_B_s that did not provide synaptic input to the CSDns (data not shown).

**Figure 6:**
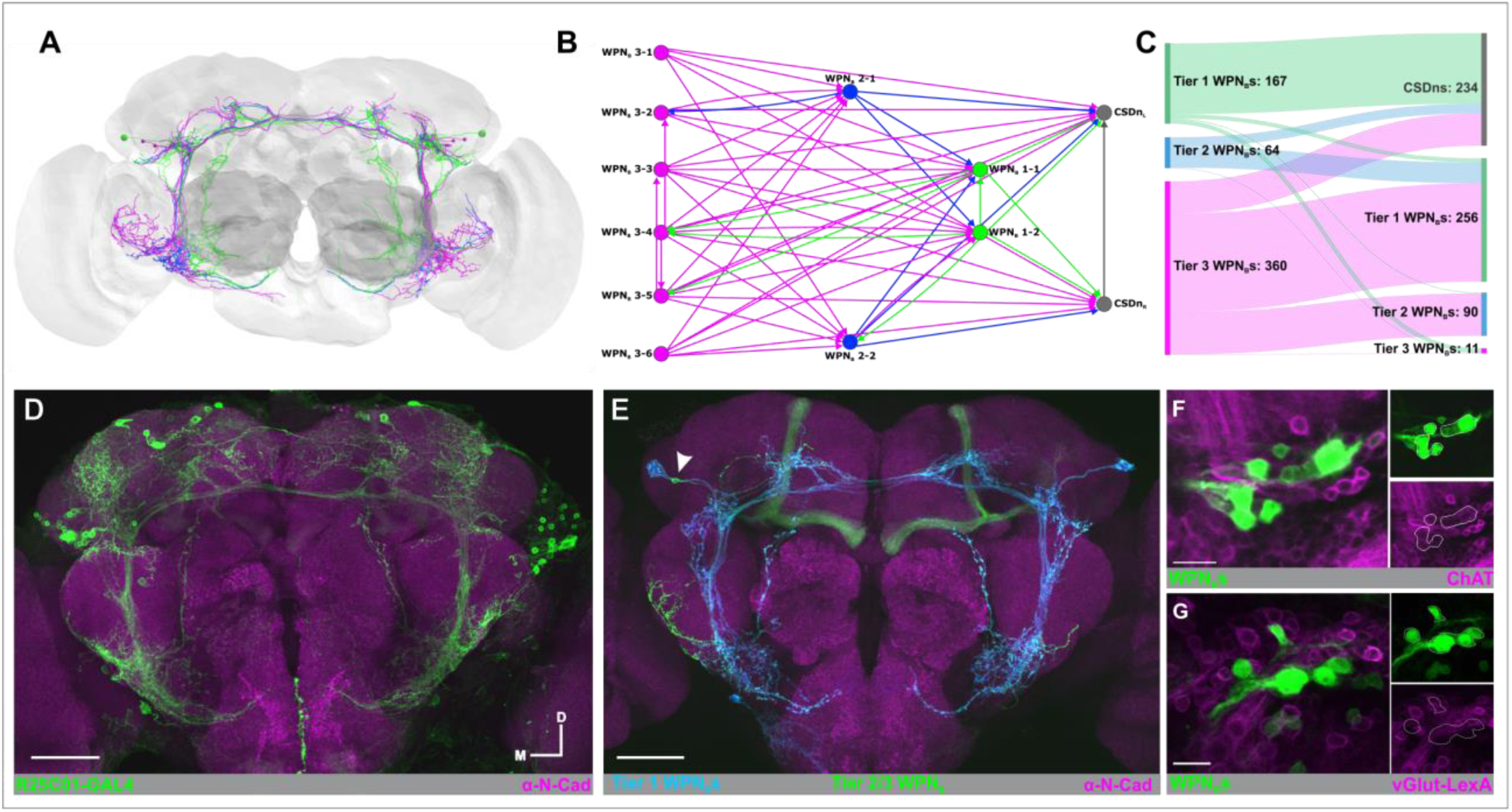
WPN_B_s provide top-down input to the CSDn. (A) EM reconstruction of 12 WPN_B_s that provide a top down input to the CSDn in the antler and portions of the protocerbrum. (B) The WPN_B_s provide input to the CSDns in a 3-tiered, feed-forward network where the Tier 1 WPN_B_s (green) are morphologically distinct from Tier 2 (blue) and Tier 3 (magenta); see also Figure 6 - figure supplement 1. (C) Tier 3 WPN_B_s provide strong input to the Tier 1 and Tier 2 WPN_B_s as well as the CSDn. (D) R25C01 driven expression of GFP (green) includes populations of WPN_B_s. (E) Multicolor FlpOut of R25C01-GAI4 highlights the expression of the two Tier 1 WPN_B_s (blue) and a Tier 2/3 WPN_B_ (green, arrow). N-Cadherin delineates neuropil (magenta). (F) Some WPN_B_s’ somata (green; R25C01-GAL4) colocalize with ChAT (magenta) Trojan-LexA::QFAD protein-trap but not vGLUT(G; magenta)Trojan-LexA::QFAD protein-trap. Scale bars D,E = 50 uM, F,G = 20 uM.

The WPN_B_s form a three-tiered feedforward network in which subsets of WPN_B_s provide synaptic input to WPN_B_s in the next tiers but the subsequent tiers provide little, if any, feedback to the WPN_B_s from which they receive input (Figure 6B,C). We further classified the WPN_B_s based on their position within this connectivity network as WPN_B_1 (two neurons), WPN_B_2 (two neurons) and WPN_B_3 (six neurons). The WPN_B_1s are morphologically distinct from WPN_B_2 and WPN_B_3 neurons as they have an additional anterodorsal projecting branch that extends from the medial saddle branch and their cell bodies are larger than the other WPN_B_s (Figure 6A, Figure 6 – figure supplement 1C). Using the Multi-Color Flp-Out technique [79], we identified a GAL4 line that is expressed by both WPN_B_1s and a subset of the WPN_B_2/3s (Figure 6D, E). We then combined this GAL4 with either a Cha^MI04508^-LexA::QFAD or a VGlut^MI04979^-LexA::QFAD protein-trap line [80] and found that many of the WPN_B_s in this GAL4 line are cholinergic, but none are glutamatergic (Figure 6F,G). The WPN_B_s have weak, non-directionality selective wind responses [72] suggesting that these neurons could be relaying mechanosensory information to the CSDns. In the process of reconstructing the WPN_B_s, we also identified an additional unilaterally projecting protocerebral neuron which provides at least 90 synapses to both CSDns in the ATL, SMP, and SLP (Figure 6 – figure supplement 2).

Finally, we identified a set of four protocerebral neurons that project into the AL where they synapse extensively upon the CSDn. These neurons have their somata along the dorsal midline of the brain, project laterally into the superior medial and intermediate protocerebra and ventrally, with one branch extending into the SEZ and another into the ipsilateral AL (Figure 7A). Based on their projections, we refer to these neurons as the “SIMPAL” (Superior Intermediate/Medial Protocerebra to Antennal Lobe) neurons. The SIMPAL neurons innervate ∼23 glomeruli before crossing the antennal commissure into the contralateral AL (Figure 7A; Figure 7 – figure supplement 1). Collectively, these extrinsic neurons provide at least 188 synapses to the left-hand CSDn (Figure 7C). We found connectivity from the SIMPAL neurons to the other, right-hand CSDn (Figure 1 – figure supplement 1B), but did not quantify synapses because it is not fully reconstructed in the AL. Interestingly, we found that the SIMPAL neurons are strongly connected to the left-hand CSDn in six glomeruli in which the ORNs respond to attractive food odors (Figure 7B; Figure 7 – figure supplement 1), in particular, DP1m, DP1l and DC1 with 10-25 synapses in each. This suggests that the influence of the SIMPAL neurons on the CSDn is non-uniform, localized and potentially within the context of food attraction. We also found that the SIMPAL neurons receive significant input from the dense ABAF LNs (Figure 7D) and most strongly in the same glomeruli in which the SIMPAL neurons provide the greatest synaptic input to the CSDn (Figure 7E). This suggests that the CSDn and the SIMPAL neurons are reciprocally connected, although any influence of the CSDn on the SIMPAL neurons would be polysynaptic via the ABAF LNs.

**Figure 7:**
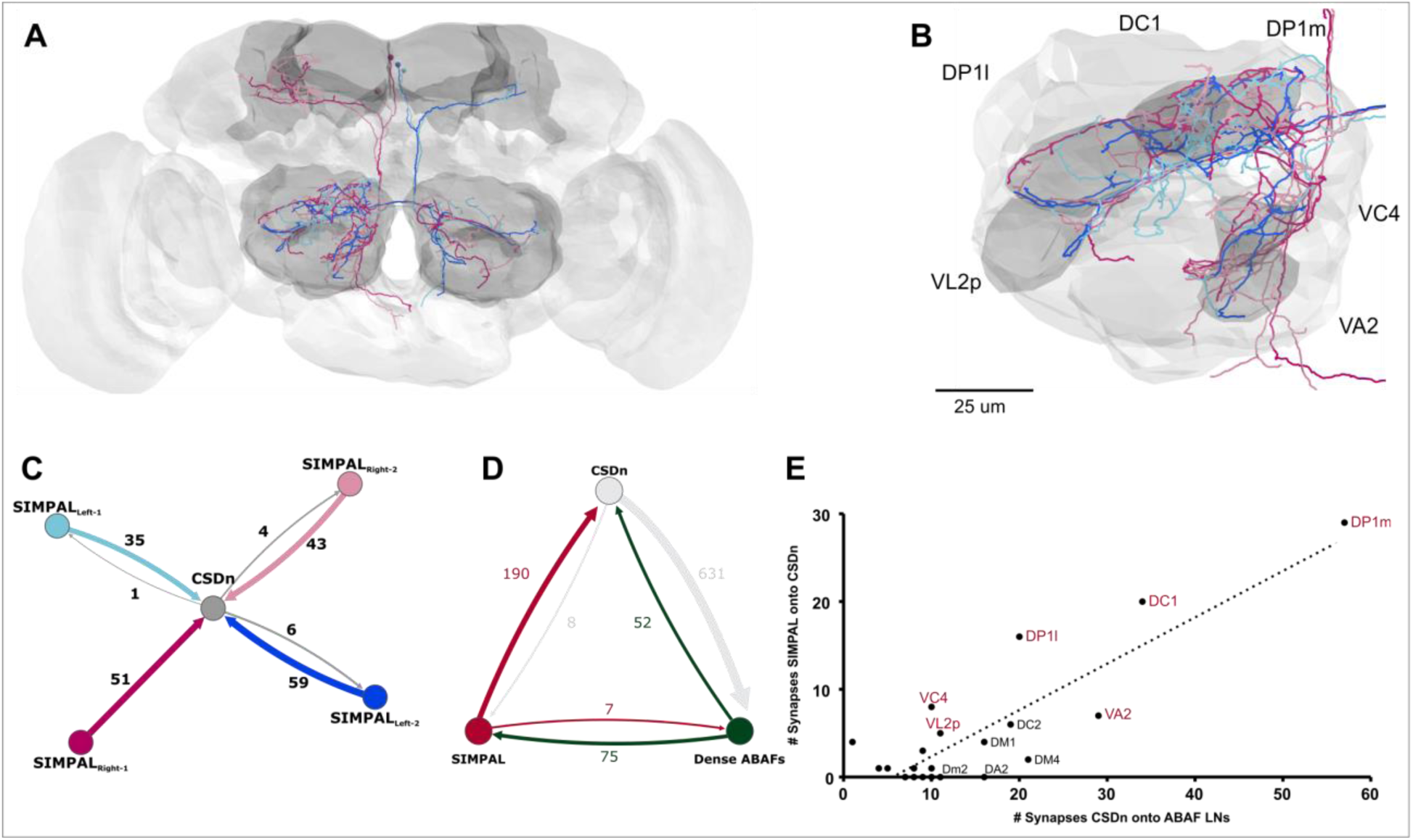
Novel Extrinsic Input from the SIMPAL neurons. (A) EM reconstruction of the four SIMPAL neurons which provide strong input (>10 synapses) to the CSDns in food odor-associated glomeruli (B) including DC1, DP1I, DP1m, VA2, VC4, and VL2p. (C) Connectivity of individual SIMPAL neurons to the CSDn is predominantly non-reciprocal. (D) The CSDns, Dense ABAF LNs, and SIMPAL neurons form a feedback loop suggesting the CSDn may influence the SIMPAL neurons polysynaptically. (E) The CSDn provides strong input to the ABAF LNs in the 6 food-odor associated glomeruli in which the SIMPAL neurons synapse upon the CSDn.

### Replication of connectivity principles across animals

While this manuscript was in preparation, a preprint describing a second dense reconstruction of a portion of a *Drosophila* central brain (called the “Hemibrain”) was made publicly available [81]. Many of the synaptic partners and connectivity relationships we found in the FAFB dataset were confirmed by the hemibrain dataset. We have included Body IDs (Figure 7-figure supplement 2) of neuron types in the Hemibrain that we describe from FAFB, although this list is not comprehensive. Thus, the broad patterns of connectivity that we describe for the CSDn are likely conserved across individual animals.

## Discussion

Heterogeneous synaptic connectivity and cell class-specific serotonin receptor expression both contribute to the complex effects of serotonin on olfactory processing and behavior. Large scale, whole brain EM datasets [82–86], can provide a comprehensive understanding of the synaptic connectivity of individual serotonergic neurons within and across networks and thus can inform predictions about the function of serotonin within discrete networks. In this study, we used a whole brain EM dataset to reconstruct a broadly projecting, identified serotonin neuron within the olfactory system of *Drosophila,* and determine its synaptic connectivity at a single-cell resolution within and between olfactory networks. Across glomeruli, the CSDn receives synaptic input from distinct combinations of olfactory neuron subpopulations, while uniformly providing synaptic input predominantly to specific subpopulations of LNs. Furthermore, the CSDn has a distinct wiring logic between the AL and LH, even establishing different connectivity relationships with the same uPN within each olfactory neuropil. Lastly, we identified neuron populations extrinsic to the olfactory system which provide strong input to the CSDn at locations within the protocerebrum and in select food-associated AL glomeruli, respectively. Population-level diversity and heterogeneity allow modulatory neurons to influence a diverse range of behaviors, often in a context-dependent manner [3-5, 8-16, 87, 88]. Our study illustrates the extent to which a single serotonergic neuron can have distinct connectivity within and across olfactory brain regions. These distinct patterns of connectivity likely allow modulatory neurons to simultaneously integrate local network information with extrinsic input to influence multiple stages of sensory processing.

### A single modulatory neuron has heterogeneous connectivity within and between sensory processing stages

The influence of serotonin and serotonergic neurons can be remarkably complex within a neural network. Within the olfactory system of vertebrate and invertebrates, serotonin can have stimulus-specific effects on odor-evoked responses or affect only a subset of a given neuron class [89–94], likely due to polysynaptic consequences of modulation and the diversity of serotonin receptors expressed in olfactory networks [45, 95–101]. For instance, serotonin differentially causes direct and polysynaptic excitation of mitral cells in the main olfactory bulb [102–104], yet direct and polysynaptic inhibition of mitral cells in the accessory olfactory bulb [105]. Furthermore, the combined release of serotonin and glutamate from dorsal raphe neurons differentially affects mitral cells and tufted cells in the main olfactory bulb, decorrelating odor-evoked responses of mitral cells and enhancing the odor-evoked responses of tufted cells [103]. This could enhance the discriminability of mitral cell odor representations while simultaneously enhancing the sensitivity of tufted cells, thus allowing simultaneous modulation of different stimulus feature representations. We found that the CSDn is differentially connected to distinct LN and PN subtypes (Figures 3 and 4), and LN and PN subtypes in *Drosophila* have different serotonin receptor expression patterns [45]. Thus, the combined complexity of cell class-specific receptor expression and heterogeneous synaptic connectivity likely underlie the seemingly variable effects of serotonin on olfactory processing.

Single serotonergic neurons can span sensory networks within the same modality and therefore have the potential to modulate different processing stages simultaneously. Recent work mapping the projection patterns of 50 serotonergic Raphe neurons demonstrated that similarly to the CSDn, a single neuron can target several olfactory areas including the main olfactory bulb, accessory olfactory bulb, and piriform cortex [19]. The degree to which the influence of a single serotonergic neuron is conserved across processing stages is unknown, however work in mice suggests that the functional roles of serotonergic neurons can differ between networks. Raphe stimulation differentially affects the odor-evoked responses of mitral and tufted cells in the main olfactory bulb [103] yet only affects spontaneous activity rather than the odor-evoked responses of single units recorded in primary piriform cortex [106]. Consistent with the idea that serotonin may play different functional roles across processing stages, we found several differences in the general connectivity rules of a single serotonergic neuron between olfactory processing stages. In the AL, the CSDn provides concentrated input to subsets of LNs (Figure 3), however CSDn input to LHLNs is far more diffuse (Figure 5). The degree to which the CSDn synapses with different populations of PNs also differs between the AL and LH (Figure 4). For instance, the CSDn is reciprocally connected to uPNs in both the AL and LH, but rarely to the same uPNs across both regions. Furthermore, the CSDn has relatively low connectivity to mPNs in the LH compared to the AL, suggesting that it preferentially targets these neurons at one processing stage. Recently it was demonstrated that local synaptic input allows the CSDn to exhibit odor invariant inhibition in the AL, yet odor and region-specific excitation in the LH [44]. This, in combination with our observation that CSDn connectivity differs between the AL and LH, suggests that single serotonergic neurons likely play different functional roles across processing stages.

### Modulatory neuron connectivity based on odor coding space

Topographical representations of stimulus features (such as stimulus identity or location) are a fundamental property of sensory systems [107–110] and overlaid upon these sensory maps are the differential projections of modulatory neurons. For instance, the density of serotonergic innervation varies between glomerular layers of the olfactory bulb [111–113] and between different glomeruli [47] and even subregions of single glomeruli in the AL [43, 114]. Although CSDn active zone density varies across glomeruli (Figure 1F; [40]), the overall demographics of CSDn postsynaptic partners were reasonably consistent across glomeruli regardless of odor tuning, with LNs being the predominant target (Figure 2A). This complements previous anatomical and physiological studies showing that GABAergic LNs synapse reciprocally with the CSDn [40], that LNs as a whole population express all five serotonin receptors [45], and the CSDn monosynaptically inhibits a population of AL LNs [42]. Here, we demonstrate the specific LN types with which the CSDn is preferentially connected, providing strong unidirectional input to densely branching ABAF LNs and reciprocal connectivity to patchy LNs (Figure 3F). Although we did not fully reconstruct the synaptic partners of the different LN subtypes, LNs synapse upon many principal neurons within the AL and support a wide range of neural computations within the AL [31-33, 35, 37-39, 62, 115]. By having different synaptic connectivity with specific LN subtypes, the CSDn may preferentially influence or actively participate in select neural computations that may be supported by these different LN types.

What then is the significance of the non-uniform glomerular innervation of the CSDn? The number of CSDn active zones within a glomerulus depends entirely upon CSDn cable length (Figure 1F), and the combination of synaptic inputs to the CSDn from ORNs, PNs, and LNs is heterogeneous across glomeruli (Figure 2B). However, the glomerulus-specific sets of neurons providing input to the CSDn are not correlated to tuning breadth alone or the density of CSDn innervation within a given glomerulus. Many odors inhibit the CSDn, with inhibition scaling with the degree to which the AL is activated [42]. Glomerulus-specific differences in input demographics suggest that the CSDn may further integrate local synaptic input within the AL in an odor-specific manner. For instance, ∼50% of the input to the CSDn in VM1 is from ORNs, compared to ∼90% of input in VM3 originating from LNs. Thus, the balance of excitation and inhibition experienced by the CSDn, and thus the influence of the CSDns on olfactory processing, likely depends in part on the odors that are currently being encountered. Alternatively, variation in input to the CSDn across glomeruli may serve to provide a broad sampling of network activity, that can be superseded in a context-dependent manner, for example, in the case of receiving input from a strongly connected partner such as the SIMPAL neurons or specific LN types.

### The CSDn as an intrinsic and extrinsic modulatory neuron

Modulatory neurons can act in either an intrinsic or extrinsic capacity [116, 117]. Intrinsic modulatory neurons receive input within the neural networks that they target and thus their influence is dependent upon recent network history. The activity of extrinsic modulatory neurons, on the other hand, is regulated by neurons outside of their target network, and thus they provide information about ongoing activity from other neural networks. In this manner, the CSDn serves as both an intrinsic and extrinsic modulatory neuron, as it has both diffuse reciprocal connectivity with the AL and LH, and receives strong, top-down input from neurons outside of these networks.

One source of top-down input to the CSDns is the WPN_B_s which form a feed-forward, hierarchical network of input (Figure 6). All three tiers within the hierarchy of the WPN_B_s provide input to the CSDns. However, tier 1 WPN_B_s integrate input from the remaining tiers and provide the greatest amount of synaptic input to the CSDns (Figure 6B, C). While the WPN_B_s are weakly activated by lateral wind input, many WPN types that show distinct tuning to wind stimuli [72]. As a population, the WPNs likely encode many features of mechanosensory input, including wind stimuli, which can be integrated with olfactory cues to locate and orient to food and mates. Each WPN_B_ tier may therefore provide information about different aspects of ongoing mechanosensory input to the antennae, such as direction, intensity, or vibration frequency. Furthermore, the positioning of their input along the main process, where the CSDn branch diameter is relatively wide [44], may allow the WPN_B_s to provide input that can spread to CSDn processes across olfactory neuropils.

The CSDn also receives strong synaptic input within the AL from four protocerebral neurons, the “SIMPAL” neurons (Figure 7). Although the role of the SIMPAL neurons is not known, they provide their greatest input to the CSDn in glomeruli that are tuned to attractive food odors such as limonene (DC1; [118]), acetic acid (DP1l; [119]), apple cider vinegar (DP1m, VA2; [120]), 2,3 butanedione (VA2; [120, 121]), ethyl lactate (VC4; [122]), and 2-oxovaleric acid (VL2p; [123, 124]). The SIMPAL neurons may affect the influence of the CSDn during the processing of food odors. Although the CSDn has almost no direct synaptic feedback to the SIMPAL neurons, they likely provide polysynaptic feedback via the ABAF LNs in these food responsive glomeruli. Nevertheless, the influence of the SIMPAL neurons upon the CSDn is likely constrained to the AL and tempered by ongoing network dynamics. In contrast, the WPN_B_s synapse upon the CSDns at multiple locations throughout the protocerebrum along the widest CSDn process, potentially allowing for greater influence over CSDn compartments.

### Concluding remarks

From the connectivity of a single serotonergic neuron, we can draw several parallels to broad organizational principles observed in larger populations of serotonergic neurons. As a population, serotonergic raphe neurons integrate input from 80 anatomically distinct areas [8, 9, 11] which allow them to respond to a diverse set of stimuli. Different populations of dorsal raphe neurons can respond to immediate events such as reward, punishment or both [18], in some cases by altering different features of their spike patterning [125]. The activity of serotonergic neurons also varies over longer time courses associated with broader physiological states [126, 127]. This complex connectivity likely supports the context-dependent effects of the dorsal raphe, for instance evoking escape behaviors under high threat conditions yet reducing movement under low threat conditions [128]. Thus, as a population, serotonergic neurons are diverse in their influence and response properties. The heterogeneity of CSDn connectivity suggests that this degree of complexity may be conserved even at the level of single serotonin neurons.

Even considering differences within and between neural networks, as well as the simultaneous integration of local and extrinsic synaptic input, the CSDns are likely more heterogeneous than their connectivity would suggest. For instance, somatodendritic and axonal expression of the serotonin re-uptake transporter by the CSDns have different effects on odor-guided behavior [41], suggesting that compartment-specific protein localization allows further functional heterogeneity. Furthermore, serotonergic neurons can also release serotonin in a paracrine manner [129, 130] potentially providing an additional form of communication. In addition, serotonergic neurons often release other transmitters, such as acetylcholine in the case of the CSDns [42] and glutamate, GABA, neuropeptides or nitric oxide in the case of serotonergic raphe neurons [18, 21, 131-133]. Thus, the synaptic influence of the CSDns likely arises from the influence of serotonin in combination with other transmitters [42]. Finally, the impact of serotonin depends upon a suite of serotonin receptors, that differ in their modes of action and binding affinity for serotonin [134, 135]. Within the AL, serotonin receptors are expressed by different principal neuron subtypes [45], adding further complexity to the influence of the CSDns. Several brain regions are densely innervated by the CSDns and each region has its own distinct network architecture. Heterogeneous, compartment-specific connectivity likely provides the CSDns with the ability to engage with the local nuances of each network while simultaneously integrating input from extrinsic sources. Given the known diversity of neurons within modulatory nuclei, it is likely that complex connectivity of individual modulatory neurons, such as the one described here, is a conserved feature of modulatory neurons across taxa.

### Methods Immunocytochemistry

Flies were raised on standard cornmeal/agar/yeast medium at 25°C and 60% humidity on a 12:12 light/dark cycle. The following fly stocks were used:

- MultiColor FlpOut (MCFO^-1^) (Bloom. #64085) and R14C11-GAL4 (Bloom. #49256) (Figure 1A)
- MB465c-split GAL4 (Bloom. #68371), UAS-Brp-short_mStraw_, UAS-GFP [55, 58] (Figure 1D)
- R25C01-GAL4 (Bloom. #49115), MultiColor FlpOut (MCFO^-1^) (Bloom. #64085), ChAT-LexA (Bloom. #60319), vGlut-LexA (Bloom. #60314), UAS-RFP, LexAop-GFP (Bloom. #32229) (Figure 6D-G)

For immunocytochemistry, brains were dissected in Drosophila external saline [136] and fixed in 4% paraformaldehyde for 30 min at 4°C. Brains were then washed in PBST (PBS with 0.5% Triton X-100), blocked for 1hr in 2% BSA (Jackson ImmunoResearch Laboratories; #001-000-162) in PBST, and incubated in primary antibodies according to Table 1. Secondary antibodies were then applied for 24 hours. Brains were then washed, ran through an ascending glycerol series (40%, 60%, and 80%), and mounted in VectaShield (Vector Labs H-1000). Images were acquired using either a 40x or 60x oil immersion lens on an Olympus FV1000 confocal microscope. Images were processed in Olympus Fluoview FV10-ASW and ImageJ.

**Table 1.**
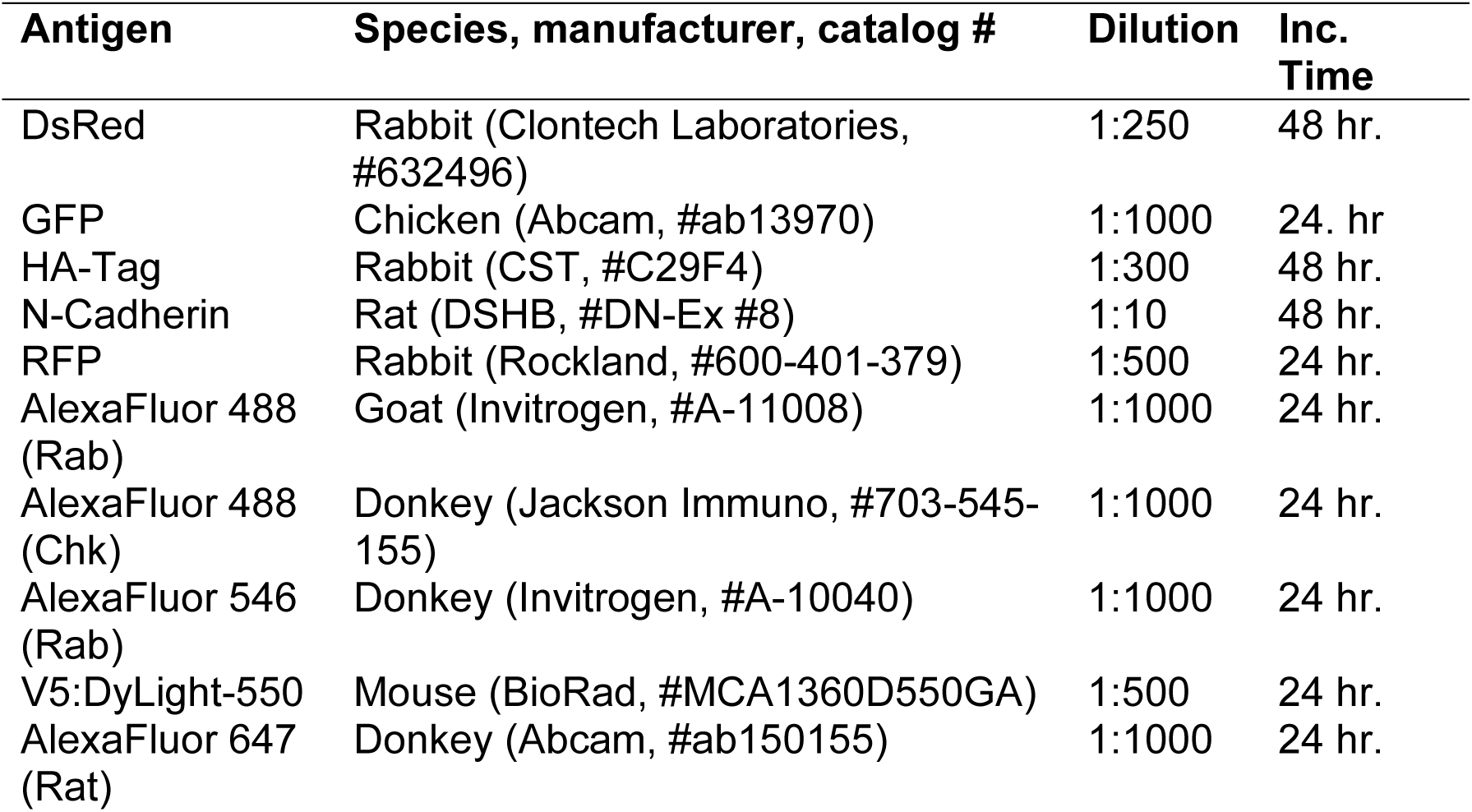
Antibodies used for Immunocytochemistry.

| Antigen | Species, manufacturer, catalog # | Dilution | Inc. Time |
| --- | --- | --- | --- |
| DsRed | Rabbit (Clontech Laboratories, #632496) | 1:250 | 48 hr. |
| GFP | Chicken (Abcam, #ab13970) | 1:1000 | 24. hr |
| HA-Tag | Rabbit (CST, #C29F4) | 1:300 | 48 hr. |
| N-Cadherin | Rat (DSHB, #DN-Ex #8) | 1:10 | 48 hr. |
| RFP | Rabbit (Rockland, #600-401-379) | 1:500 | 24 hr. |
| AlexaFluor 488 (Rab) | Goat (Invitrogen, #A-11008) | 1:1000 | 24 hr. |
| AlexaFluor 488 (Chk) | Donkey (Jackson Immuno, #703-545-155) | 1:1000 | 24 hr. |
| AlexaFluor 546 (Rab) | Donkey (Invitrogen, #A-10040) | 1:1000 | 24 hr. |
| V5:DyLight-550 | Mouse (BioRad, #MCA1360D550GA) | 1:500 | 24 hr. |
| AlexaFluor 647 (Rat) | Donkey (Abcam, #ab150155) | 1:1000 | 24 hr. |

### EM Dataset and Neuron Reconstruction

The whole female adult fly brain (FAFB) Electron Microscopy dataset was previously generated as described in [25]. The dataset is available for download at https://fafb.catmaid.virtualflybrain.org/, https://www.temca2data.org. Neuron reconstructions were “traced” manually in FAFB using CATMAID (http://www.catmaid.org) as previously described [137, 138]. In brief, reconstructions were generated by following the confines of a neuron’s cellular membrane and adding place markers (nodes). For a given neuron, all nodes are connected. Thus, users were able to reconstruct the morphology of a neuron throughout the brain volume as well as annotate synapses. A synapse was identified by the presence of 1) t-bars 2) vesicle cloud, 3) synaptic cleft, and 4) post-synaptic density [25]. It should be noted that gap junctions cannot be visualized within the EM dataset and bulk release sites are not readily evident. All reconstructions were verified by a second, experienced tracer [25, 138]. Some neurons were also reconstructed in part using an automated segmentation version of the FAFB dataset [139]. These auto-traced neuron fragments were concatenated by expert users in a separate instance of FAFB and then imported and merged with previously traced fragments in the actively traced FAFB dataset using the Python tool “FAFBseg” (https://github.com/flyconnectome/fafbseg-py) and verified for accuracy. Finally, the PNs and LH neurons used in this analysis were published elsewhere [25, 48–50, 123].

### CSDn Reconstruction

The left-hand CSD neuron (i.e. soma in the fly’s left hemisphere) was originally identified based on the unique omega-shaped projection pattern of its primary process that spans the protocerebrum. The CSDn’s identity was later confirmed using NBLAST [51] to query the reconstructed skeleton against a dataset of light microscopy images of single neurons [52, 140]. The primary process of the CSDn was reconstructed from soma to ipsilateral protocerebrum to contralateral protocerebrum, to contralateral AL. Arborizations from the primary process into the lateral horn and antennal lobes were also reconstructed “to completion” using methods described in [25]. For all CSDn branches traced, all presynaptic and postsynaptic sites, as well as pre- and post-synaptic partners were marked. In total, 23,328,903 nm of CSDn cable was reconstructed with 2,885 presynaptic sites and 4,141 postsynaptic sites marked. It should be noted that an incomplete reconstruction of the CSDn was previously published in [44]. It should also be noted that we identified and partially reconstructed the right-hand CSDn (soma in the fly’s right hemisphere) (Figure 1 – supplement 1B) and observed two synapses between the two CSDns.

### Reconstruction of CSDn partners in 9 glomeruli

We chose nine glomeruli to reconstruct all CSDn synaptic partners in the contralateral AL based on the lifetime sparseness of the glomerulus as well as the number of pre and post-synaptic sites of the CSDn (Figure 1F; Figure 2 – figure supplement 1). To reconstruct synaptic partners in specific glomeruli, the completed reconstruction of the CSDn was filtered in CATMAID so that only CSDn branches and synapses within a specific glomerulus were visible, using glomerulus volume meshes generated in [50]. All neurons synapsing upon the CSDn or receiving synaptic input from the CSDn were then reconstructed in a given glomerulus towards the neuron’s primary neurite (i.e. backbone) and ceased when the reconstruction was sufficient to identify the neuron as a PN, LN, ORN, or other neuron type based on its soma location and/or projection pattern. Short neuron fragments that could not be reconstructed to backbone due to ambiguity of projections were deemed “orphans” and were excluded from analyses. Reconstructions were reviewed from starting synapse to the backbone (primary neurite) by a second expert tracer and then queried using NBLAST as needed.

### LN reconstruction and classification

In the process of reconstructing CSDn synaptic partners in the 9 select glomeruli, we found that the CSDn is synaptically connected with 84 LNs. LNs were reconstructed to the extent that, at a minimum, allowed them to be morphologically characterized and synapses with the CSDn were marked. 2 Patchy LNs, 2 dense ABAF, and 1 sparse ABAF were reconstructed from soma to backbone into the finer processes of each glomerulus to synapse dense regions.

To compare LNs within and across morphological types, KNN analyses were performed between all of a neuron branch points and their 1^st^, 3^rd^, and 8^th^ nearest branch points. This generated a distribution of the distance between neighboring branch points for each neuron so that distributions could be compared between neurons. Similar branch point distributions therefore indicated that individual LNs likely belong to the same morphological grouping. Neurons that had more branch points were randomly subsampled so that an equal number of branch points were compared across neurons. Analyses scripts and related figures were generated in R using analysis packages within the natverse (http://natverse.org/) [141].

### CSDn active zone distribution in the LH

To determine the distribution of active zones in the lateral horn, CSDn active zones were labeled using Brp-short_mstraw_ as described in [40]. Confocal z-stacks of the lateral horn were acquired and an “intensity distribution analysis” was performed using Matlab as previously described in [114]. In brief, the Brp puncta intensity was averaged and normalized across 10×10 bins and plotted as heatmaps.

### Neuronal skeleton data analysis

#### Input/Output Index

The “Input/Output Index” of presynaptic and postsynaptic sites of the CSDn was calculated by generating a list of presynaptic connectors and postsynaptic connectors within a given neuropil or glomerulus volume using PyMaid (https://github.com/schlegelp/pymaid). Volumes used were generated in [50]. For each neuropil, the “Input/Output Index” was calculated as (# presynaptic sites / (# presynaptic + # postsynaptic sites).

#### Other Analyses

Synapse fractions, connectivity graphs, and connectivity matrices were generated using CATMAID. The volume of each neuropil mesh, branch length per glomerulus and the number of synapses per glomerulus was calculated using PyMaid (https://github.com/schlegelp/pymaid) and analyzed using GraphPad Prism.

### Synapse fractions PCA

PCAs of synapse fractions and associated analyses were performed in R using the *ggfortify, ggplot, and factorextra* libraries (https://rpkgs.datanovia.com/factoextra/index.html). K-means clustering was done to determine if points clustered in the PCA, using PC-scores. (https://www.rdocumentation.org/packages/stats/versions/3.6.2/topics/kmeans) [142]. To determine the number of clusters for the k-means, the silhouette method [143] was performed using a k-max of 4, as PC1-PC4 explained more of the variance than would be expected if the variance were equally distributed across all 9 PCs as determined by the scree plot. K-means clustering coloring was then applied to the PCA.

To determine if the variability of upstream and downstream partners differed, the Euclidean distance between each downstream point with all other downstream points were measured in a pairwise manner. The Euclidean distance was determined using PC1-PC4 as explained above (Figure 2 – figure supplement 5B’’) for the downstream partners of the 9 glomeruli and then the upstream partners of the 9 glomeruli. Statistical differences in downstream versus upstream distances were determined using a Student’s t-test.

### Imaging processing and analysis

Images of the EM dataset, connectivity graphs, and reconstructions were acquired and exported from CATMAID. The rendered skeleton of the CSDn shown in Figure 1C was generated using Blender (https://www.blender.org/) and the CATMAID-to-Blender plugin (https://github.com/schlegelp/CATMAID-to-Blender). All figures were organized using CorelDrawX9.

## Data availability

Neuron reconstructions generated by our group will be uploaded to the open-access website https://catmaid-fafb.virtualflybrain.org upon publication.

## Acknowledgments

We are greatly indebted to the FAFB tracing community for their helpful insights and contributions of neuron tracings, Tom Kazimiers and Andrew Champion for CATMAID development, Peter Li for development of the autosegmented instance of the FAFB dataset, and Eric Perlman for making the autosegmented dataset available in CATMAID. In particular, we would like to thank the Cambridge Drosophila Connectomics group, especially Greg Jefferis, Marta Costa, Philipp Schlegel, Alex Bates, Ruairi Roberts, Robert Turnbull, Lisa Marin, and Nik Drummond for use of hundreds of neuron reconstructions, assistance with custom analyses, and facilitating access to the autosegmented FAFB dataset. We would also like to thank Feng Li, Mert Erginkaya, Johann Schor, Jeremy Johnson, Shrey Patel, and Nick Sweet for contributions of neuron tracings or review; Masayoshi Ito for identifying the R25C01-GAL4 line as expressing the WPN_B_s; Jay Milam for assistance with collecting confocal scans of the WPN_B_ transmitter colocalization; Tessa Cessario for fly care; Tyler Sizemore for assistance with recombining fly stocks; Quentin Gaudry for providing the UAS-RFP, LexAop-GFP;vGlut-LexA line, Tim Mosca for providing the UAS-Brp-short_mStraw_, UAS-GFP and David Krantz for providing the hsFLP;;MCFO-1 line. Finally, we would like to thank Kevin Daly, Tim Mosca, and Tyler Sizemore for feedback on the manuscript and Kevin Daly, Gary Marsat, and Keshav Ramachandra for insight on data analysis. This work was supported by an NIH DC 016293 and a USAFOSR FA9550-17-1-0117 to AMD, the HHMI Janelia Research Campus Visiting Scientist Program project to AMD and KEC, and a Wellcome Trust Collaborative Award (203261/Z/16/Z) and NIH RF1 MH120679 01 award to DDB. The HHMI supported the generation and hosting of the FAFB dataset.

---

**Figure 1 - figure supplement 1:**
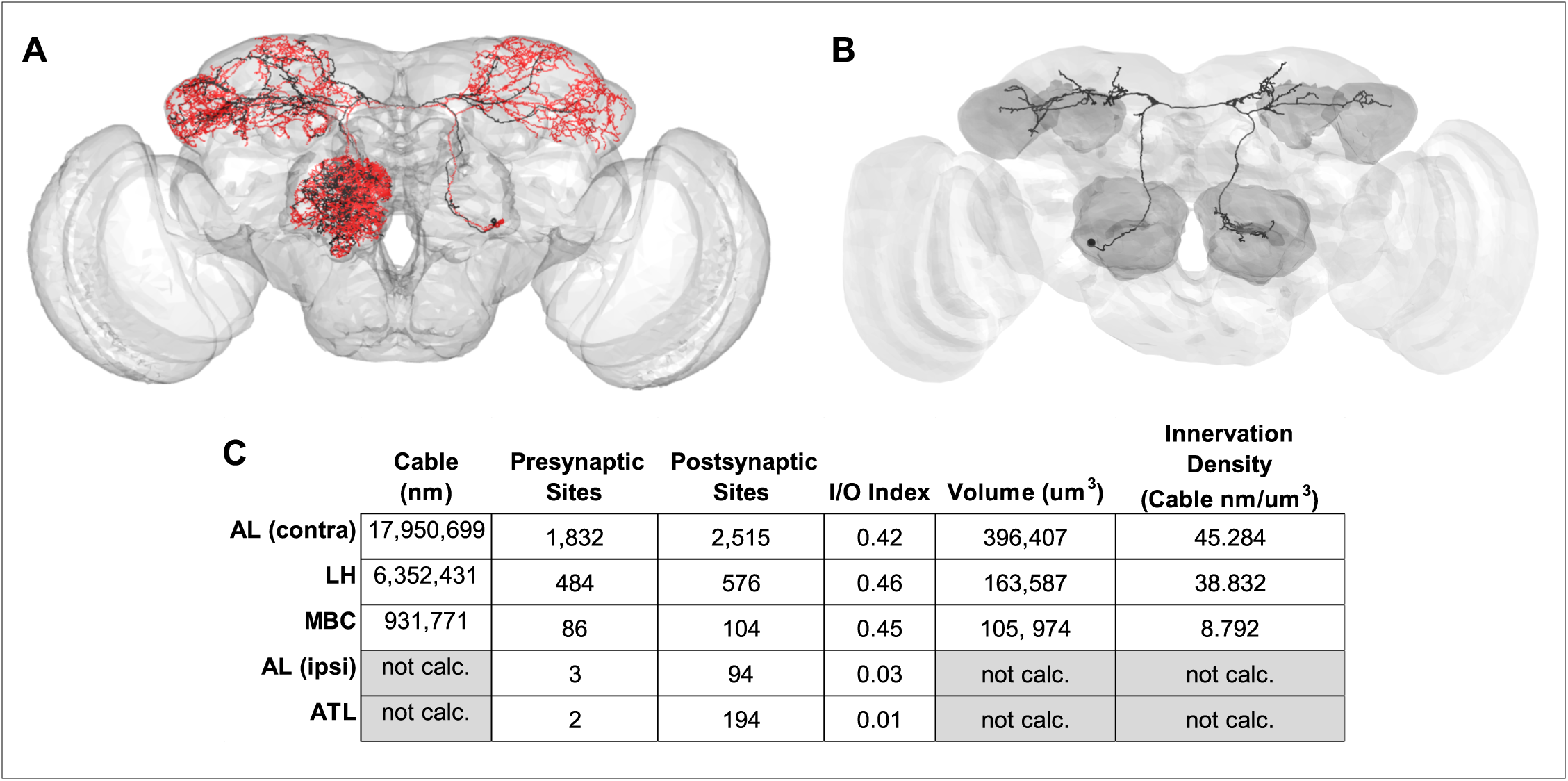
NBLAST and stats of the left-hand CSDn and EM reconstruction of the right-hand CSDn. (A) Overlay of the (left-hand) CSDn reconstruction (black) with skeletonized CSDn from light microscopy image dataset via NBLAST (red). Similarity score = 0.716 (where 1 equals perfect alignment with dataset). (B) Partial Reconstruction of the right-hand CSDn. (C) Broad stats of the CSDn across neuropil, including metrics used to make the ‘‘heat maps” in Figure 1C.

**Figure 1 - figure supplement 2:**
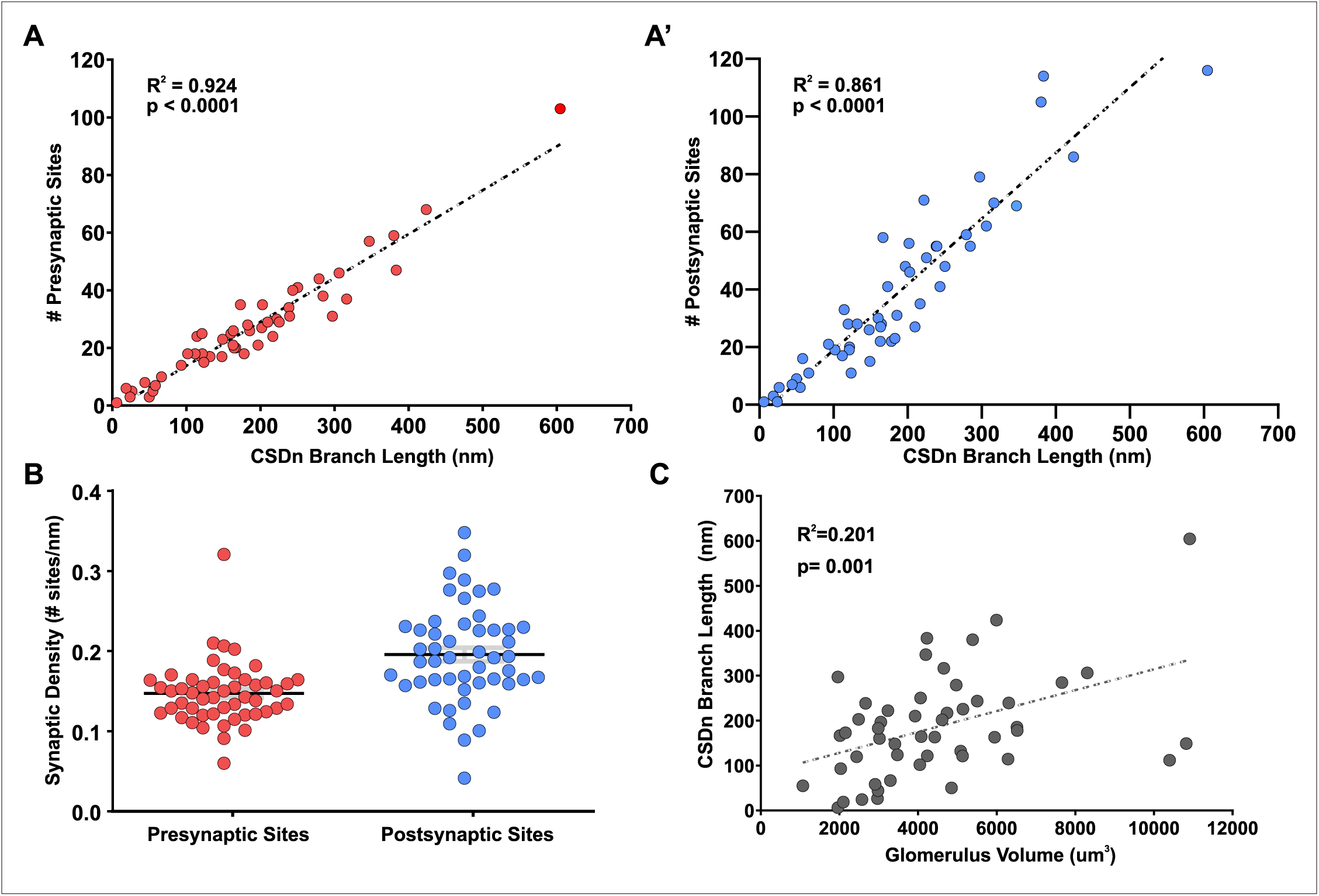
CSDn AL stats. CSDn branch length in glomeruli is highly correlated to (A) CSDn presynaptic sites (R^2^=0.924, p < 0.0001) and (A’) number of CSDn postsynaptic sites (R^2^=0.861, p < 0.0001). (B) CSDn presynaptic density (red) (# sites/nm branch length) and postsynaptic density (blue) are fairly consistent across glomeruli (COV = 26.26% and 30.85%, respectively), although postsynaptic is more distributed (p < 0.005, Levene’s test for homogeneity of variance). (C) CSDn branch length per glomerulus weakly correlates to glomerular volume (R^2^=0.201, p = 0.001).

**Figure 2 - Figure Supplement 1:**
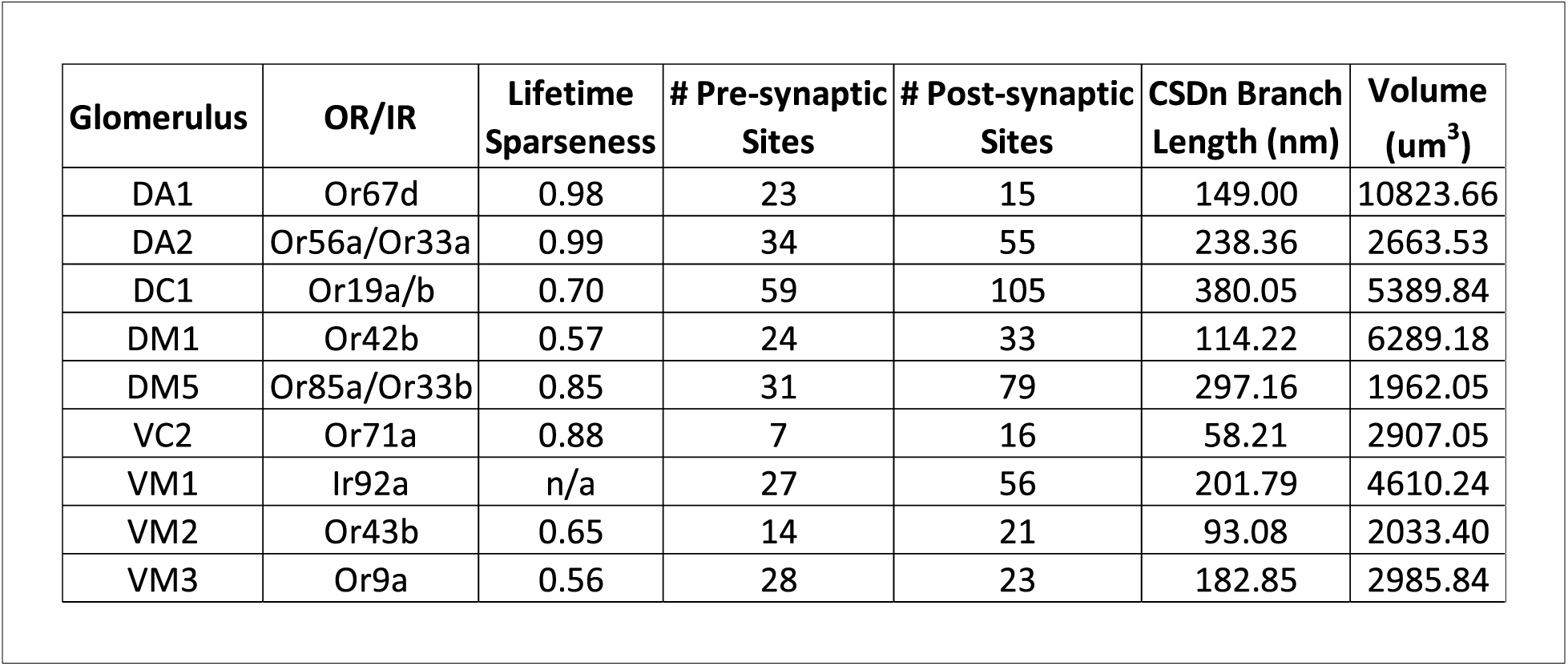
Data for glomeruli in which CSDn synaptic partners were reconstructed. Glomeruli were chosen for reconstruction based on the glomerulus’s lifetime sparseness, number of pre and postsynaptic sites, CSDn branch length and glomerulus volume to get an array of different combinations of features.

**Figure 2 - figure supplement 2:**
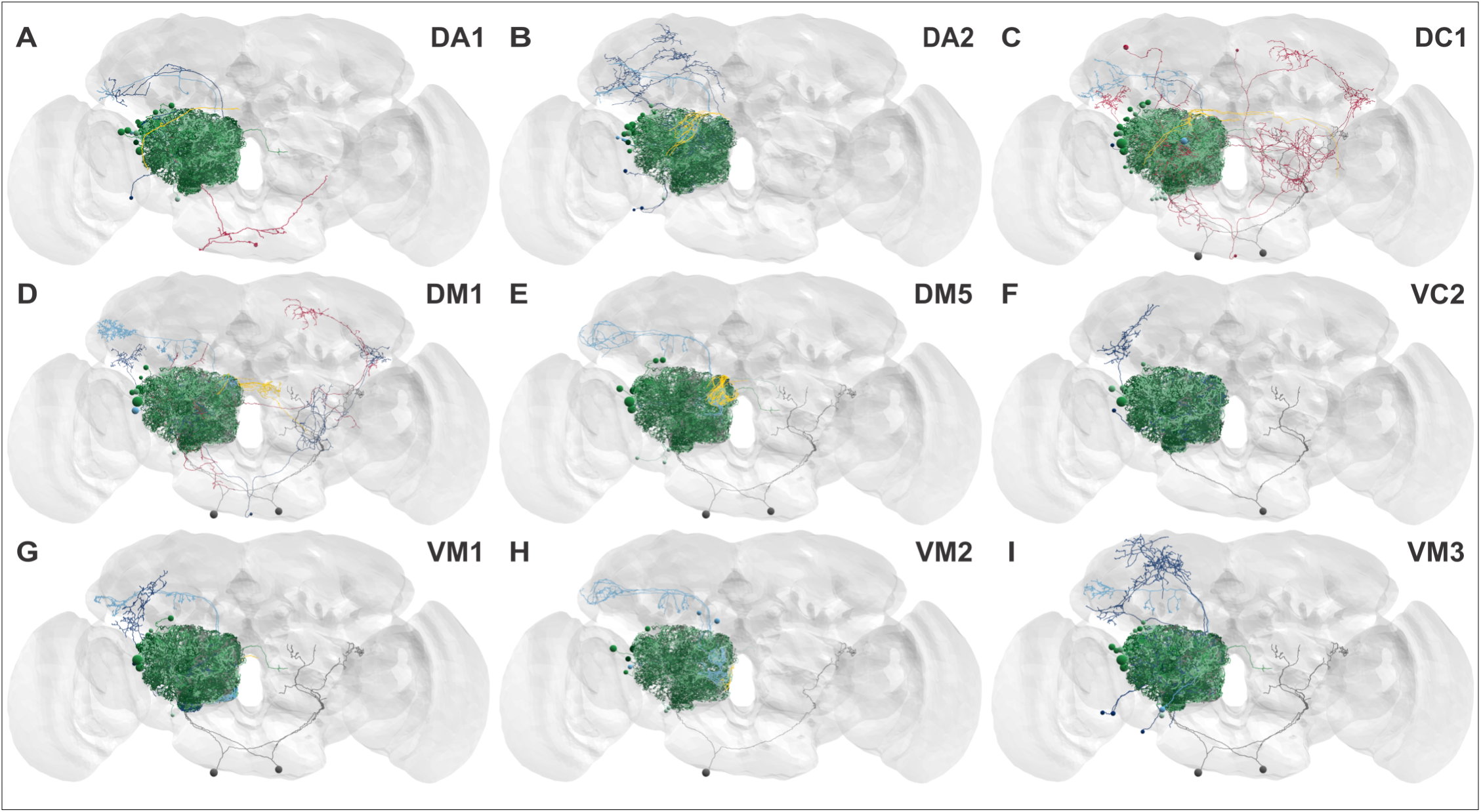
Downstream synaptic partners of the CSDn in 9 glomeruli within the AL. EM reconstructions of all neurons that receive input from the CSDn across the 9 glomeruli in the AL: DA1 (A), DA2 (B), DC1 (C), DM1 (D), DM5 (E), VC2 (F), VM1 (G), VM2 (H), and VM3 (I). Neuron were grouped as ORNs (yellow), LNs (green shades), PNs (blue shades), and extrinsic neurons (red).

**Figure 2 - figure supplement 3:**
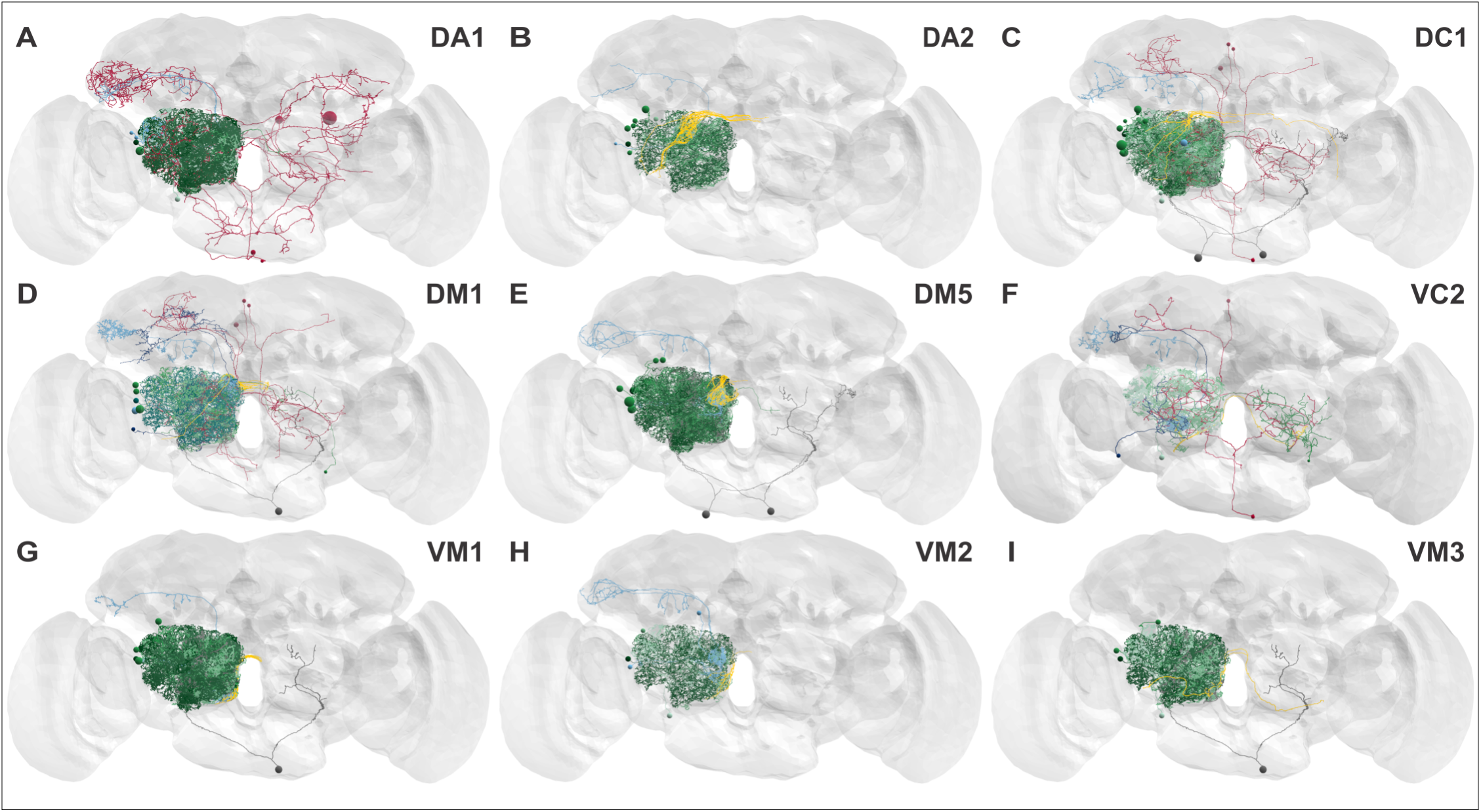
Upstream synaptic partners of the CSDn in 9 glomeruli within the AL. EM reconstructions of all neurons that provide input to the CSDn across the 9 glomeruli in the AL: DA1 (A), DA2 (B), DC1 (C), DM1 (D), DM5 (E), VC2 (F), VM1 (G), VM2 (H), and VM3 (I). Neuron were grouped as ORNs (yellow), LNs (green shades), PNs (blue shades), and extrinsic neurons (red).

**Figure 2 - figure supplement 4:**
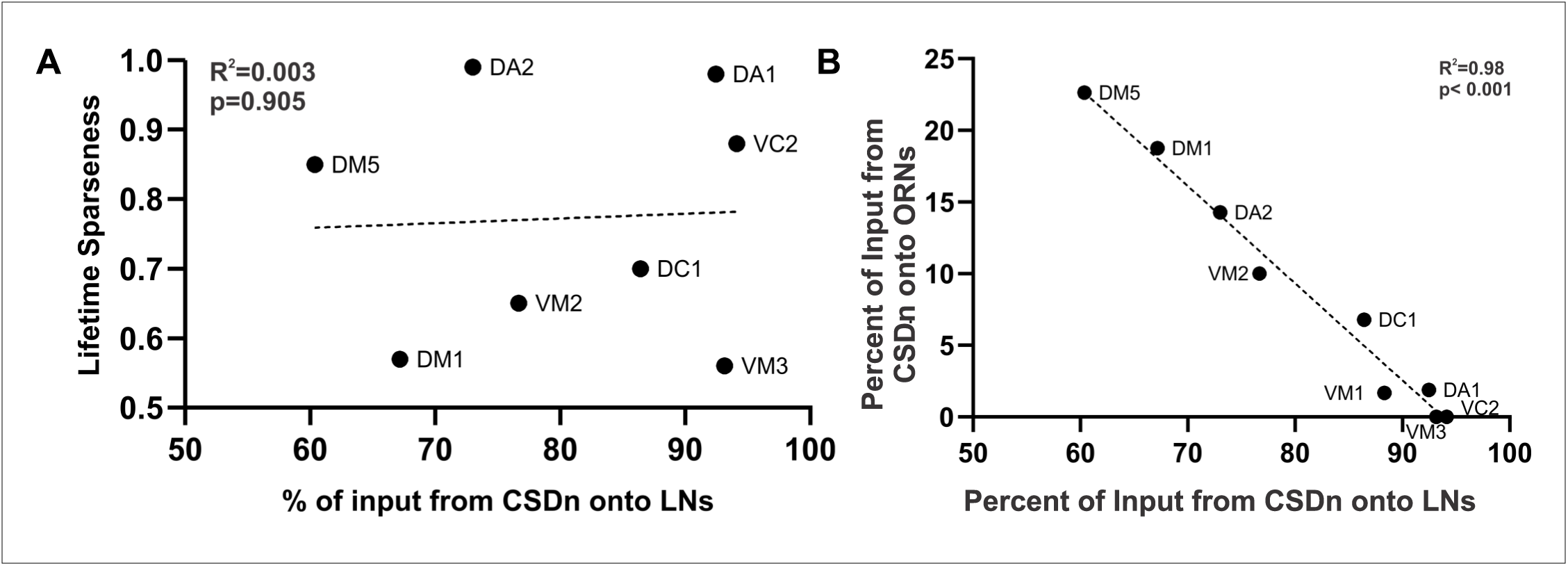
CSDn connectivity relationships across glomeruli. (A) Percent of input from the CSDn onto LNs does not correlated with lifetime sparseness of a glomerulus (R^2^=0.003). VM1’s lifetime sparseness value is unknown, thus is excluded. (B) Fraction of input from the CSDn onto LNs is inversely correlated to the percent of input from the CSDn onto ORNs.

**Figure 2 - figure supplement 5:**
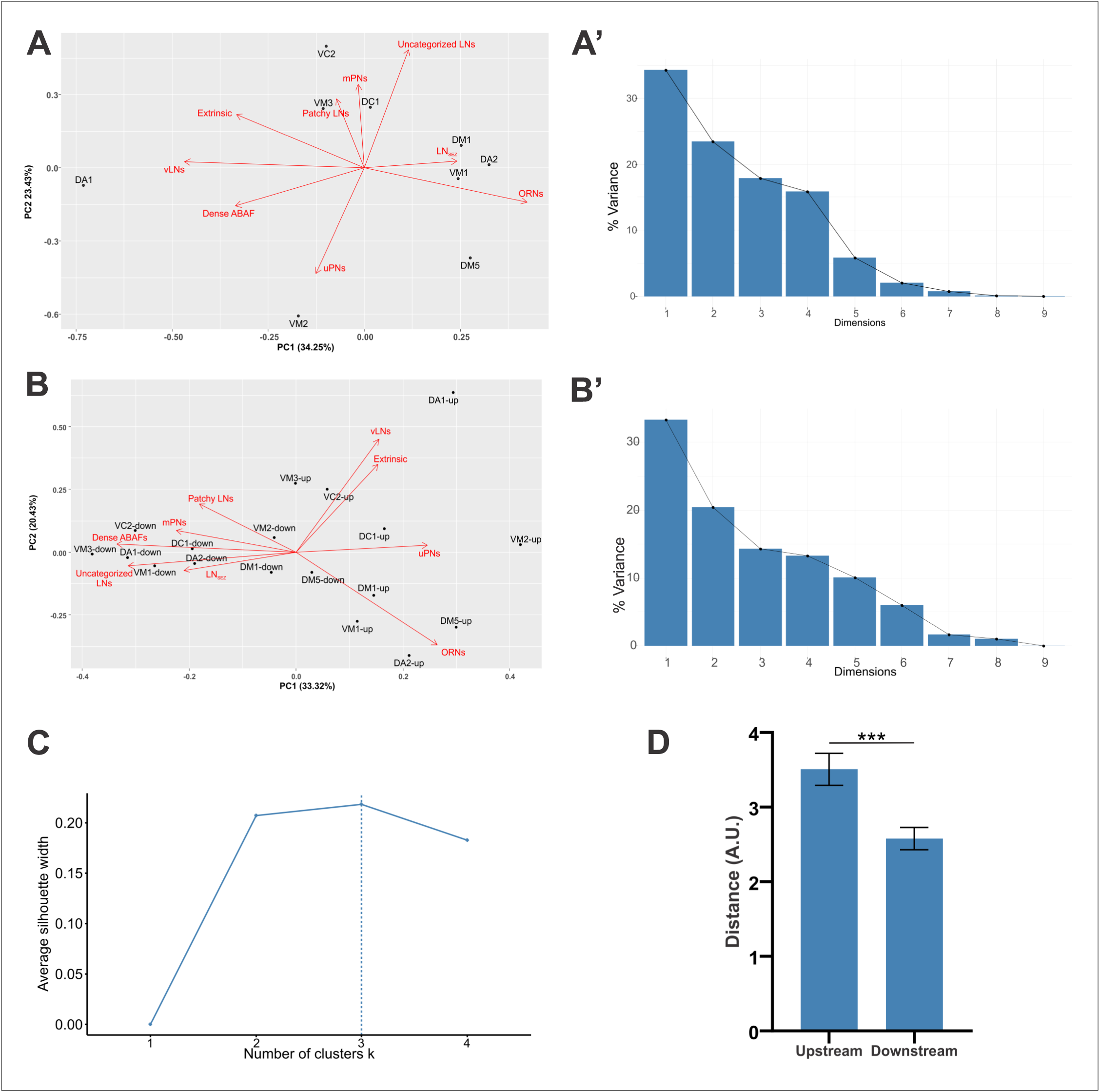
PCA. (A) PCA of upstream synapse fractions shows that glomeruli do not cluster in PC space based on eigenvectors. (A’) Variance is explained by the first 4 principal components. (B) PCAof up and downstream synaptic partners from Figure 2C with eigenvectors shown. (B’) Scree plot used to determine which principal components explain the most variance. (C) Silhouette method used to determine the optimal number of clusters for K-means clustering in Figure 2C. (D) The mean distance between downstream points in the PCA (B’) is significantly different from the mean distance between upstream points (p=0.0007, Student’s t-test).

**Figure 3 - figure supplement 1:**
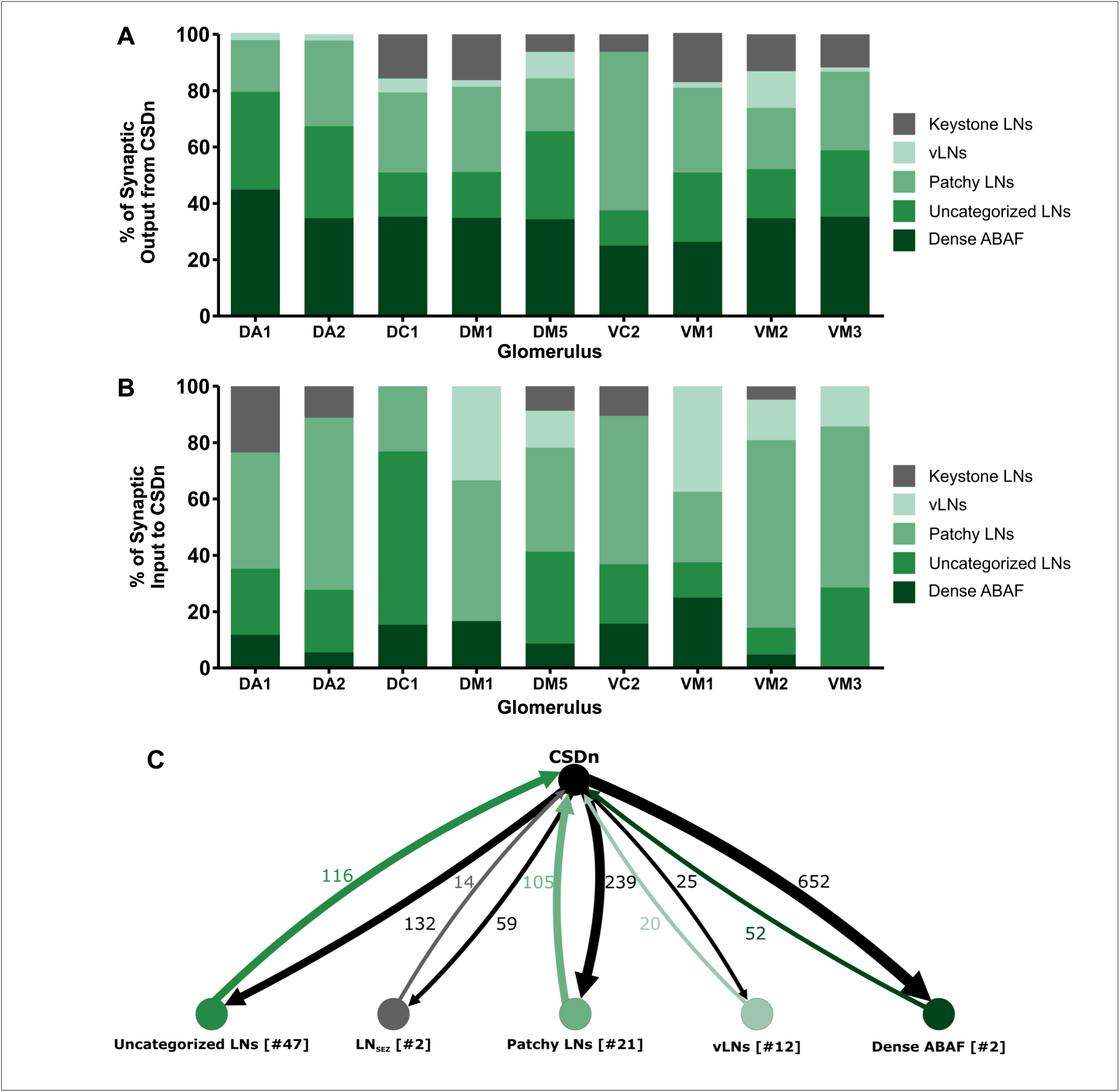
CSDn input from and to LN subtypes varies across glomeruli. (A) Percent of synaptic input from the CSDn onto LN subtypes. The CSDn provides input to most LN subtypes across all 9 glomeruli, except for Keystone-llke LNs. Dense ABAFs and Patchy LNs appear to be the main LN type which the CSDn targets regardless of glomerulus identity. (B) The CSDn receives far more of its LN synaptic input from Patchy LNs across 9 glomeruli. (C) Number of synapses of the CSDn with each subtype of LN across the AL.

**Figure 3 - figure supplement 2:**
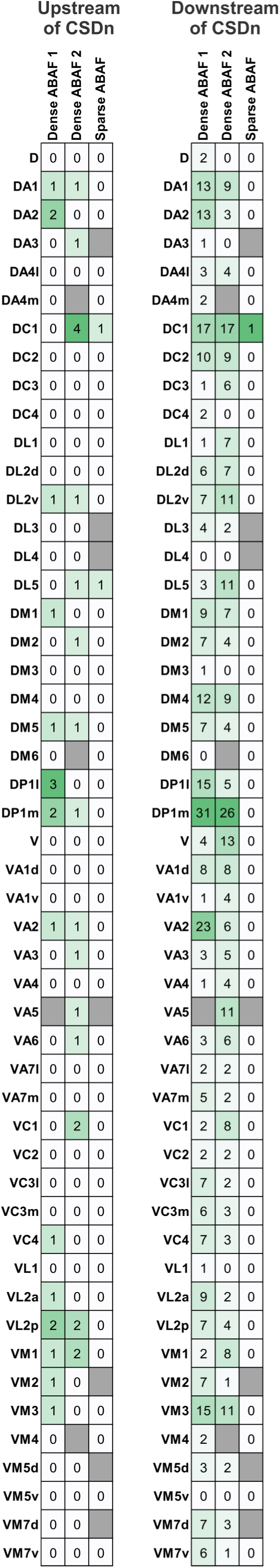
ABAF LN connectivity with the CSDn across glomeruli. Number of synapses to and from the CSDn with the two dense ABAFs and the sparse ABAF in each glomeruli. Left = from ABAF onto CSDn, Right = CSDn onto ABAF. All ventral-posterior glomeruli and the VC5 glomerulus are excluded. Gray boxes = glomeruli where the LN does not innervate (dictated by having no branchpoints in the glomerulus).

**Figure 3 - figure supplement 3:**
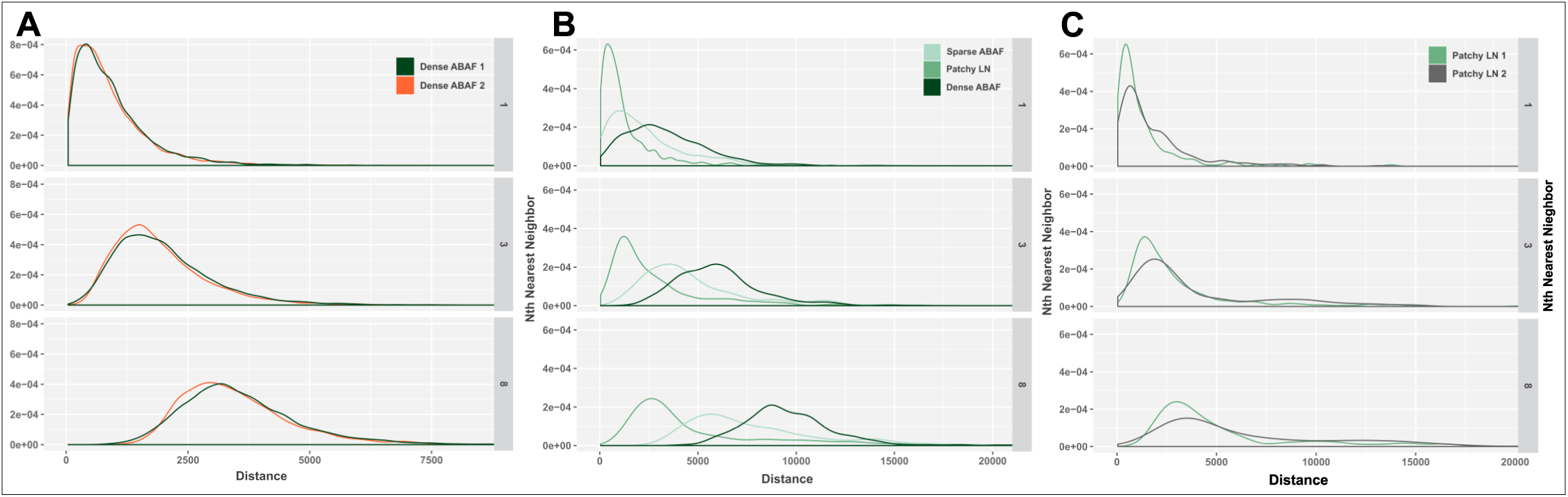
K-Nearest Neighbor analyses. (A) Distribution of KNN analysis showing that the distance of the nearest neighboring branch points of the two Dense ABAFs is consistent. Thus, they belong to the same morphological class. (B) KNN showing that Sparse ABAF and Dense ABAF LNs belong to two different morphological sub-classes. The Patchy LN is included as an outgroup. (C) KNN distribution showing that two patchy LNs belong to the same morphological class. Plots show distance from the 1st (top), 3rd (middle), and 8th (bottom), nearest neighboring branch points.

**Figure 4 - Figure supplement 1:**
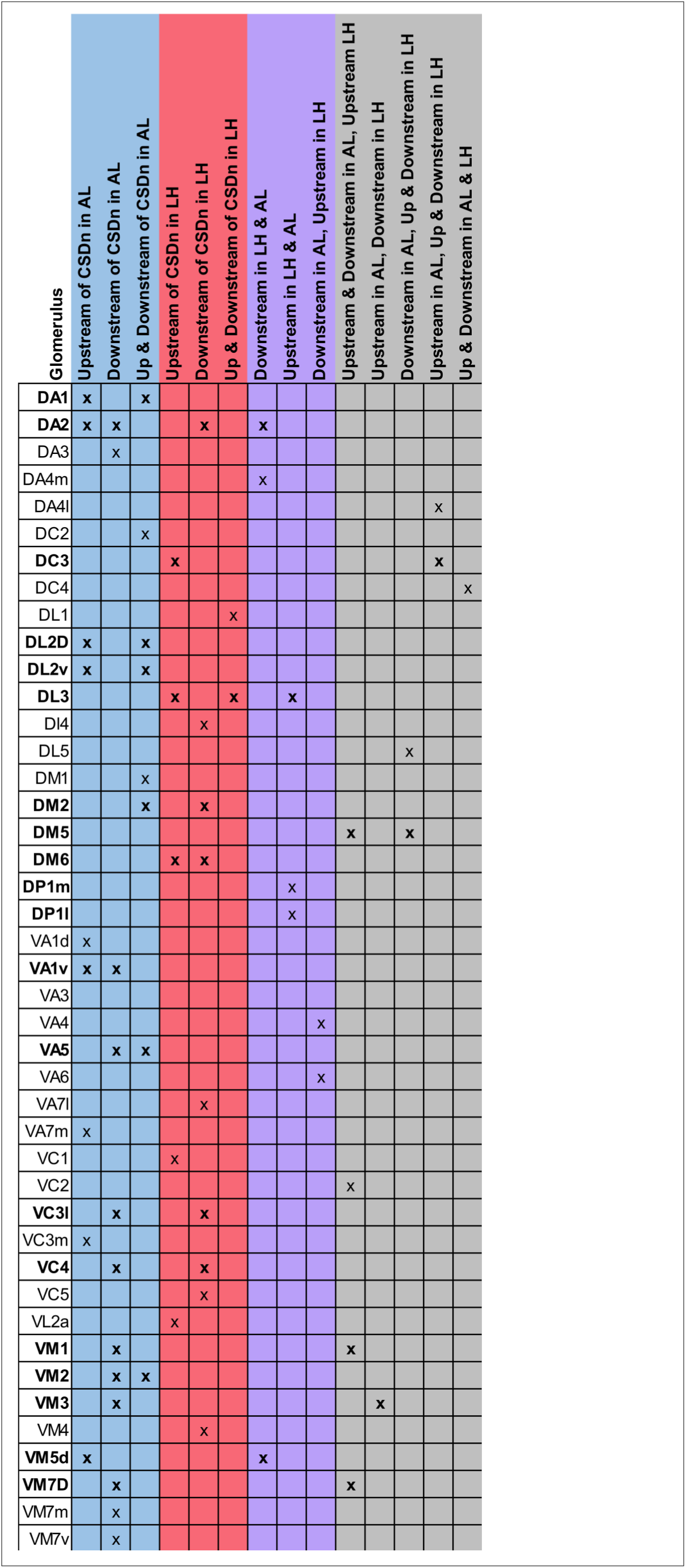
CSDn Connectivity with uPNs in the AL and LH. Summary table of the types of connectivity individual uPNs have with the CSDn across the AL and LH. Blue = AL only, Red = LH only, Purple = AL & LH, Gray = various combinations of upstream and downstream connectivity. Bolded names represent glomeruli that have uPNs with several different combinations of synaptic connectivity with the CSDn.

**Figure 4 - figure supplement 2:**
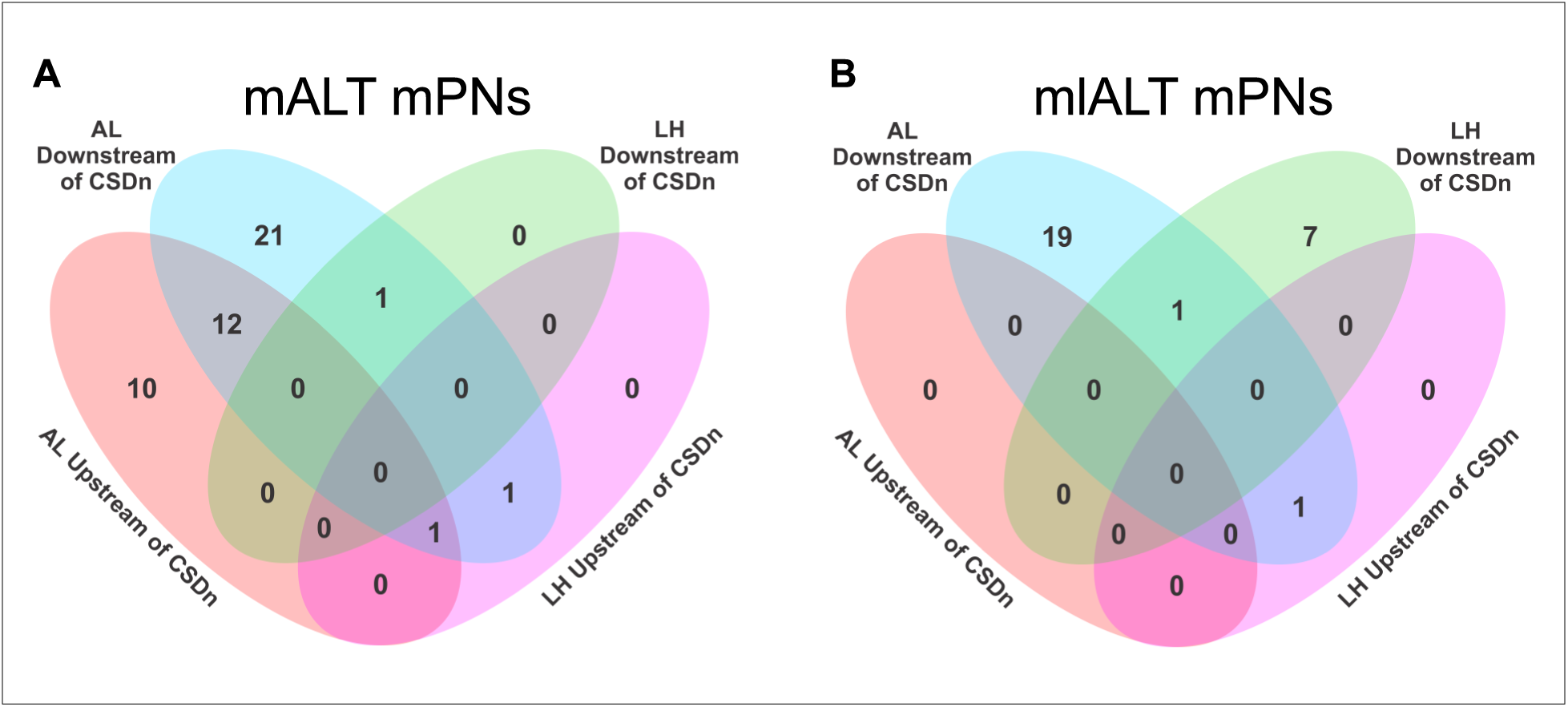
CSDn Connectivity with mPNs in the AL and LH. Representation of the number of individual (A) mALT mPNs and (B) mlALT mPNs that receive synaptic input from the CSDn (i.e. downstream), provide synaptic input to the CSDn (i.e. upstream), or both across the AL and LH.

**Figure 5 - figure supplement 1:**
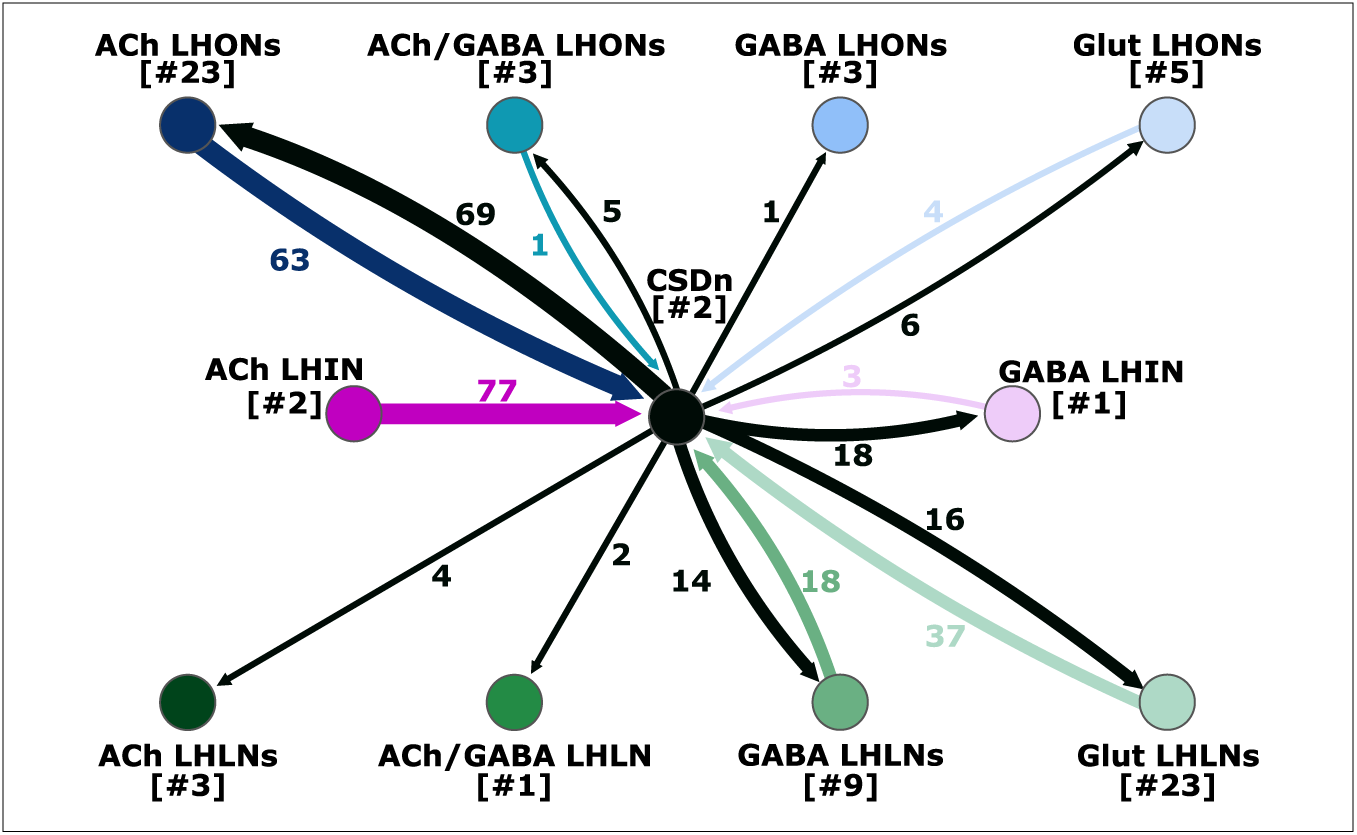
CSDn’s Connectivity with lateral horn neurons based on transmitter content, including weak connectivity.

**Figure 6 - figure supplement 1:**
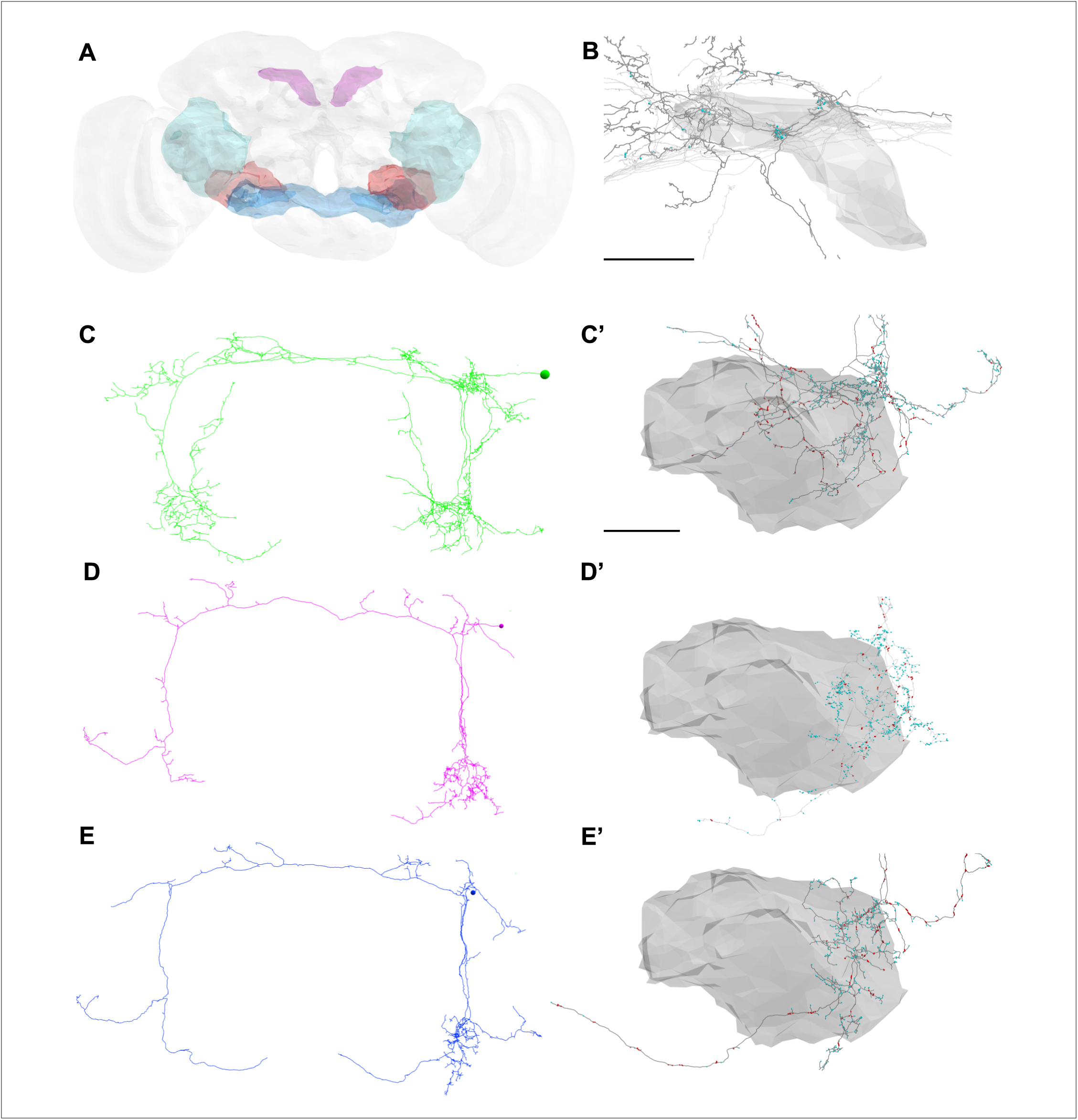
Three tiers of WPN_B_s. (A) The WPN_B_s innervate the highlighted neuropil: ATL (magenta), AVLP (cyan), Saddle (blue), and Wedge (red). (B) WPN_B_s (light gray) provide input to the CSDn (dark gray) in the ATL. CSDn postsynaptic sites markers are shown in cyan. EM reconstructions of a Tier 1 WPN_B_ (C; green), Tler2 WPN_B_ (D; magenta) and Tier 3 WPN_B_ (E; blue). The Tier 1 WPN_B_s have two morphologically distinct branches projecting antero-dorsally and larger somata than the Tier 2 and Tier 3 WPN_B_s. All three tiers of WPN_B_s have dendritic regions in the wedge (blue dots; C’,D’, E’). Red dots along each skeleton indicate presynaptic sites. Scale Bars = 25 uM.

**Figure 6 - figure supplement 2:**
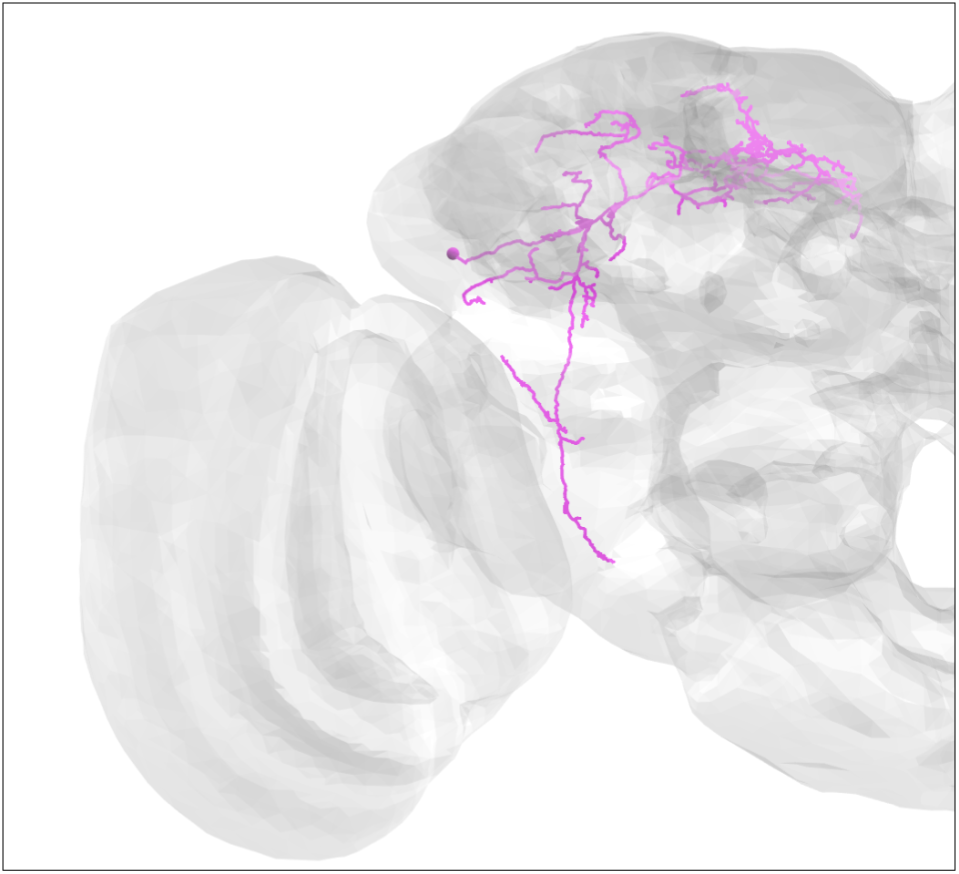
EM reconstruction of a previously undescribed protocerebral neuron that provides strong input (at least 90 synapses) to the CSDns in the Superior Medial Protocerebrum, Superior Lateral Protocerebrum, and Antler.

**Figure 7 - figure supplement 1:**
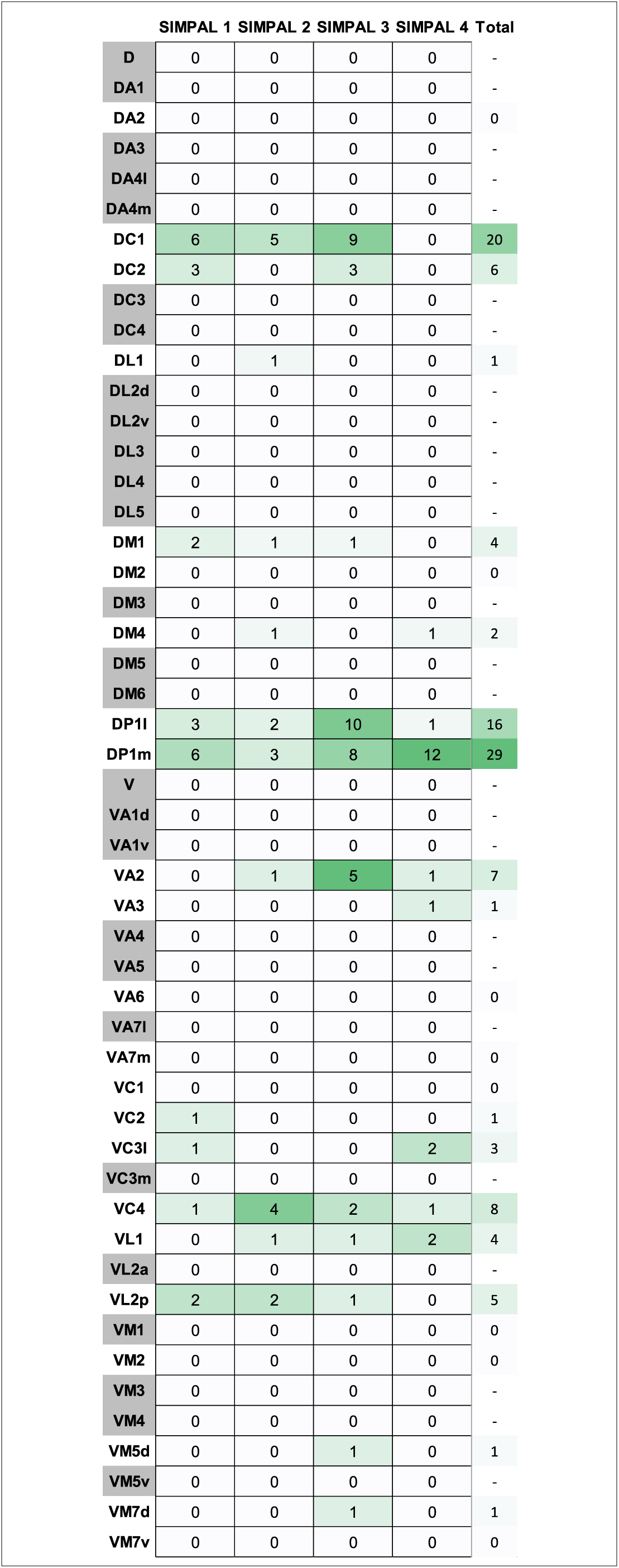
Number of synapses from the four SIMPAL neurons onto the CSDn in each glomerulus in the AL. Glomeruli shaded gray are those which the SIMPAL neurons do not innervate.

**Figure 7 - figure supplement 2:**
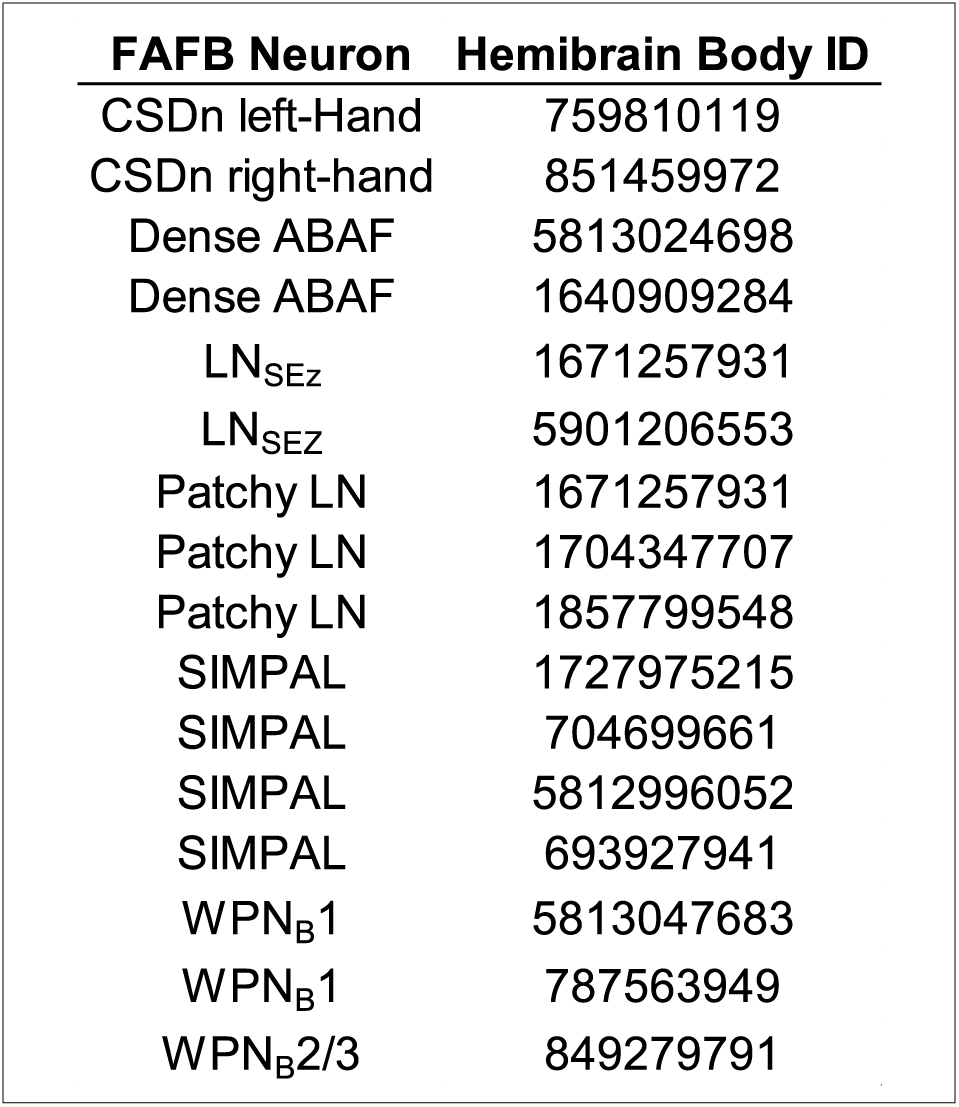
Several key synaptic partners of the CSDn we reconstructed in the FAFB dataset (left) were also identified in the Hemibraln dataset based on morphology (right).

## References

1. Marder, E., Neuromodulation of neuronal circuits: back to the future. Neuron, 2012. 76(1): p. 1–11.

2. Nadim, F. and D. Bucher, Neuromodulation of neurons and synapses. Curr Opin Neurobiol, 2014. 29: p. 48–56.

3. Morales, M. and E.B. Margolis, Ventral tegmental area: cellular heterogeneity, connectivity and behaviour. Nat Rev Neurosci, 2017. 18(2): p. 73–85.

4. Okaty, B.W., K.G. Commons, and S.M. Dymecki, Embracing diversity in the 5-HT neuronal system. Nat Rev Neurosci, 2019. 20(7): p. 397–424.

5. Schwarz, L.A. and L. Luo, Organization of the locus coeruleus-norepinephrine system. Curr Biol, 2015. 25(21): p. R1051–R1056.

6. Halliday, G.M., et al., Distribution of monoamine-synthesizing neurons in the human medulla oblongata. J Comp Neurol, 1988. 273(3): p. 301–17.

7. Ishimura, K., et al., Quantitative analysis of the distribution of serotonin-immunoreactive cell bodies in the mouse brain. Neurosci Lett, 1988. 91(3): p. 265–70.

8. Pollak Dorocic, I., et al., A whole-brain atlas of inputs to serotonergic neurons of the dorsal and median raphe nuclei. Neuron, 2014. 83(3): p. 663–78.

9. Weissbourd, B., et al., Presynaptic partners of dorsal raphe serotonergic and GABAergic neurons. Neuron, 2014. 83(3): p. 645–62.

10. Jensen, P., et al., Redefining the serotonergic system by genetic lineage. Nat Neurosci, 2008. 11(4): p. 417–9.

11. Ogawa, S.K., et al., Organization of monosynaptic inputs to the serotonin and dopamine neuromodulatory systems. Cell Rep, 2014. 8(4): p. 1105–18.

12. Okaty, B.W., et al., Multi-Scale Molecular Deconstruction of the Serotonin Neuron System. Neuron, 2015. 88(4): p. 774–91.

13. Bang, S.J., et al., Projections and interconnections of genetically defined serotonin neurons in mice. Eur J Neurosci, 2012. 35(1): p. 85–96.

14. O’Hearn, E. and M.E. Molliver, Organization of raphe-cortical projections in rat: a quantitative retrograde study. Brain Res Bull, 1984. 13(6): p. 709–26.

15. Wylie, C.J., et al., Distinct transcriptomes define rostral and caudal serotonin neurons. J Neurosci, 2010. 30(2): p. 670–84.

16. Andrade, R. and S. Haj-Dahmane, Serotonin neuron diversity in the dorsal raphe. ACS Chem Neurosci, 2013. 4(1): p. 22–5.

17. Commons, K.G., Two major network domains in the dorsal raphe nucleus. J Comp Neurol, 2015. 523(10): p. 1488–504.

18. Ren, J., et al., Anatomically Defined and Functionally Distinct Dorsal Raphe Serotonin Sub-systems. Cell, 2018. 175(2): p. 472–487 e20.

19. Ren, J., et al., Single-cell transcriptomes and whole-brain projections of serotonin neurons in the mouse dorsal and median raphe nuclei. Elife, 2019. 8.

20. Fernandez, S.P., et al., Multiscale single-cell analysis reveals unique phenotypes of raphe 5-HT neurons projecting to the forebrain. Brain Struct Funct, 2016. 221(8): p. 4007–4025.

21. Huang, K.W., et al., Molecular and anatomical organization of the dorsal raphe nucleus. Elife, 2019. 8.

22. Katz, P.S. and P.D. Quinlan, The importance of identified neurons in gastropod molluscs to neuroscience. Curr Opin Neurobiol, 2019. 56: p. 1–7.

23. Dacks, A.M., T.A. Christensen, and J.G. Hildebrand, Phylogeny of a serotonin-immunoreactive neuron in the primary olfactory center of the insect brain. J Comp Neurol, 2006. 498(6): p. 727–46.

24. Roy, B., et al., Metamorphosis of an identified serotonergic neuron in the Drosophila olfactory system. Neural Dev, 2007. 2: p. 20.

25. Zheng, Z., et al., A Complete Electron Microscopy Volume of the Brain of Adult Drosophila melanogaster. Cell, 2018. 174(3): p. 730–743 e22.

26. Groschner, L.N. and G. Miesenbock, Mechanisms of Sensory Discrimination: Insights from Drosophila Olfaction. Annu Rev Biophys, 2019. 48: p. 209–229.

27. Liang, L. and L. Luo, The olfactory circuit of the fruit fly Drosophila melanogaster. Sci China Life Sci, 2010. 53(4): p. 472–84.

28. Wilson, R.I., Early olfactory processing in Drosophila: mechanisms and principles. Annu Rev Neurosci, 2013. 36: p. 217–41.

29. Grabe, V. and S. Sachse, Fundamental principles of the olfactory code. Biosystems, 2018. 164: p. 94–101.

30. Chou, Y.H., et al., Diversity and wiring variability of olfactory local interneurons in the Drosophila antennal lobe. Nat Neurosci, 2010. 13(4): p. 439–49.

31. Hong, E.J. and R.I. Wilson, Simultaneous encoding of odors by channels with diverse sensitivity to inhibition. Neuron, 2015. 85(3): p. 573–89.

32. Liu, W.W. and R.I. Wilson, Glutamate is an inhibitory neurotransmitter in the Drosophila olfactory system. Proc Natl Acad Sci U S A, 2013. 110(25): p. 10294–9.

33. Nagel, K.I., E.J. Hong, and R.I. Wilson, Synaptic and circuit mechanisms promoting broadband transmission of olfactory stimulus dynamics. Nat Neurosci, 2015. 18(1): p. 56–65.

34. Olsen, S.R., V. Bhandawat, and R.I. Wilson, Divisive normalization in olfactory population codes. Neuron, 2010. 66(2): p. 287–99.

35. Olsen, S.R. and R.I. Wilson, Lateral presynaptic inhibition mediates gain control in an olfactory circuit. Nature, 2008. 452(7190): p. 956-60.

36. Seki, Y., et al., Physiological and morphological characterization of local interneurons in the Drosophila antennal lobe. J Neurophysiol, 2010. 104(2): p. 1007–19.

37. Yaksi, E. and R.I. Wilson, Electrical coupling between olfactory glomeruli. Neuron, 2010. 67(6): p. 1034–47.

38. Ignell, R., et al., Presynaptic peptidergic modulation of olfactory receptor neurons in Drosophila. Proc Natl Acad Sci U S A, 2009. 106(31): p. 13070–5.

39. Root, C.M., et al., A presynaptic gain control mechanism fine-tunes olfactory behavior. Neuron, 2008. 59(2): p. 311–21.

40. Coates, K.E., et al., Identified serotonergic modulatory neurons have heterogeneous synaptic connectivity within the olfactory system of Drosophila. J Neurosci, 2017.

41. Kasture, A.S., et al., Distinct contribution of axonal and somatodendritic serotonin transporters in drosophila olfaction. Neuropharmacology, 2019. 161: p. 107564.

42. Zhang, X. and Q. Gaudry, Functional integration of a serotonergic neuron in the Drosophila antennal lobe. Elife, 2016. 5.

43. Sun, X.J., L.P. Tolbert, and J.G. Hildebrand, Ramification pattern and ultrastructural characteristics of the serotonin-immunoreactive neuron in the antennal lobe of the moth Manduca sexta: a laser scanning confocal and electron microscopic study. J Comp Neurol, 1993. 338(1): p. 5–16.

44. Zhang, X., et al., Local synaptic inputs support opposing, network-specific odor representations in a widely projecting modulatory neuron. Elife, 2019. 8.

45. Sizemore, T.R. and A.M. Dacks, Serotonergic Modulation Differentially Targets Distinct Network Elements within the Antennal Lobe of Drosophila melanogaster. Sci Rep, 2016. 6: p. 37119.

46. Xu, L., et al., A Single Pair of Serotonergic Neurons Counteracts Serotonergic Inhibition of Ethanol Attraction in Drosophila. PLoS One, 2016. 11(12): p. e0167518.

47. Singh, A.P., et al., Sensory neuron-derived eph regulates glomerular arbors and modulatory function of a central serotonergic neuron. PLoS Genet, 2013. 9(4): p. e1003452.

48. Dolan, M.J., et al., Neurogenetic dissection of the Drosophila lateral horn reveals major outputs, diverse behavioural functions, and interactions with the mushroom body. Elife, 2019. 8.

49. Frechter, S., et al., Functional and anatomical specificity in a higher olfactory centre. Elife, 2019. 8.

50. Bates, A.S., et al., Complete connectomic reconstruction of olfactory projection neurons in the fly brain. BioRxiv, 2020.

51. Costa, M., et al., *NBLAST: Rapid,* Sensitive Comparison of Neuronal Structure and Construction of Neuron Family Databases. Neuron, 2016. 91(2): p. 293–311.

52. Chiang, A.S., et al., Three-dimensional reconstruction of brain-wide wiring networks in Drosophila at single-cell resolution. Curr Biol, 2011. 21(1): p. 1–11.

53. Jeanne, J.M., M. Fisek, and R.I. Wilson, The Organization of Projections from Olfactory Glomeruli onto Higher-Order Neurons. Neuron, 2018. 98(6): p. 1198–1213 e6.

54. Jeanne, J.M. and R.I. Wilson, Convergence, Divergence, and Reconvergence in a Feedforward Network Improves Neural Speed and Accuracy. Neuron, 2015. 88(5): p. 1014–1026.

55. Fouquet, W., et al., Maturation of active zone assembly by Drosophila Bruchpilot. J Cell Biol, 2009. 186(1): p. 129–45.

56. Owald, D., et al., A Syd-1 homologue regulates pre- and postsynaptic maturation in Drosophila. J Cell Biol, 2010. 188(4): p. 565–79.

57. Schmid, A., et al., Activity-dependent site-specific changes of glutamate receptor composition in vivo. Nat Neurosci, 2008. 11(6): p. 659–66.

58. Mosca, T.J. and L. Luo, Synaptic organization of the Drosophila antennal lobe and its regulation by the Teneurins. Elife, 2014. 3: p. e03726.

59. Grabe, V., et al., Elucidating the Neuronal Architecture of Olfactory Glomeruli in the Drosophila Antennal Lobe. Cell Rep, 2016. 16(12): p. 3401–3413.

60. Carlsson, M.A., et al., Multiple neuropeptides in the Drosophila antennal lobe suggest complex modulatory circuits. J Comp Neurol, 2010. 518(16): p. 3359–80.

61. Das, A., et al., Identification and analysis of a glutamatergic local interneuron lineage in the adult Drosophila olfactory system. Neural Syst Circuits, 2011. 1(1): p. 4.

62. Wilson, R.I. and G. Laurent, Role of GABAergic inhibition in shaping odor-evoked spatiotemporal patterns in the Drosophila antennal lobe. J Neurosci, 2005. 25(40): p. 9069–79.

63. Berck, M.E., et al., The wiring diagram of a glomerular olfactory system. Elife, 2016. 5.

64. Otopalik, A.G., et al., Sloppy morphological tuning in identified neurons of the crustacean stomatogastric ganglion. Elife, 2017. 6.

65. Otopalik, A.G., et al., When complex neuronal structures may not matter. Elife, 2017. 6.

66. Tobin, W.F., R.I. Wilson, and W.A. Lee, Wiring variations that enable and constrain neural computation in a sensory microcircuit. Elife, 2017. 6.

67. Marin, E.C., et al., Connectomics analysis reveals first, second, and third order thermosensory and hygrosensory neurons in the adult Drosophila brain. BioRxiv, 2020.

68. Liang, L., et al., GABAergic projection neurons route selective olfactory inputs to specific higher-order neurons. Neuron, 2013. 79(5): p. 917–31.

69. Parnas, M., et al., Odor discrimination in Drosophila: from neural population codes to behavior. Neuron, 2013. 79(5): p. 932–44.

70. Strutz, A., et al., Decoding odor quality and intensity in the Drosophila brain. Elife, 2014. 3: p. e04147.

71. Caron, S.J., et al., Random convergence of olfactory inputs in the Drosophila mushroom body. Nature, 2013. 497(7447): p. 113-7.

72. Suver, M.P., et al., Encoding of Wind Direction by Central Neurons in Drosophila. Neuron, 2019. 102(4): p. 828–842 e7.

73. Ito, M., et al., Systematic analysis of neural projections reveals clonal composition of the Drosophila brain. Curr Biol, 2013. 23(8): p. 644–55.

74. Kamikouchi, A., et al., The neural basis of Drosophila gravity-sensing and hearing. Nature, 2009. 458(7235): p. 165-71.

75. Lai, J.S., et al., Auditory circuit in the Drosophila brain. Proc Natl Acad Sci U S A, 2012. 109(7): p. 2607–12.

76. Matsuo, E., et al., Organization of projection neurons and local neurons of the primary auditory center in the fruit fly Drosophila melanogaster. J Comp Neurol, 2016. 524(6): p. 1099–164.

77. Patella, P. and R.I. Wilson, Functional Maps of Mechanosensory Features in the Drosophila Brain. Curr Biol, 2018. 28(8): p. 1189–1203 e5.

78. Vaughan, A.G., et al., Neural pathways for the detection and discrimination of conspecific song in D. melanogaster. Curr Biol, 2014. 24(10): p. 1039–49.

79. Nern, A., B.D. Pfeiffer, and G.M. Rubin, Optimized tools for multicolor stochastic labeling reveal diverse stereotyped cell arrangements in the fly visual system. Proc Natl Acad Sci U S A, 2015. 112(22): p. E2967–76.

80. Diao, F., et al., Plug-and-play genetic access to drosophila cell types using exchangeable exon cassettes. Cell Rep, 2015. 10(8): p. 1410–21.

81. Xu, C.S., et al., A Connectome of the Adult Drosophila Central Brain. BioRxiv, 2020.

82. Bates, A.S., et al., Neuronal cell types in the fly: single-cell anatomy meets single-cell genomics. Curr Opin Neurobiol, 2019. 56: p. 125–134.

83. Schlegel, P., M. Costa, and G.S. Jefferis, Learning from connectomics on the fly. Curr Opin Insect Sci, 2017. 24: p. 96–105.

84. Cook, S.J., et al., Whole-animal connectomes of both Caenorhabditis elegans sexes. Nature, 2019. 571(7763): p. 63-71.

85. Meinertzhagen, I.A., Of what use is connectomics? A personal perspective on the Drosophila connectome. J Exp Biol, 2018. 221(Pt 10).

86. Ohyama, T., et al., A multilevel multimodal circuit enhances action selection in Drosophila. Nature, 2015. 520(7549): p. 633-9.

87. Beier, K.T., et al., Circuit Architecture of VTA Dopamine Neurons Revealed by Systematic Input-Output Mapping. Cell, 2015. 162(3): p. 622–34.

88. Schwarz, L.A., et al., Viral-genetic tracing of the input-output organization of a central noradrenaline circuit. Nature, 2015. 524(7563): p. 88-92.

89. Dacks, A.M., T.A. Christensen, and J.G. Hildebrand, Modulation of olfactory information processing in the antennal lobe of Manduca sexta by serotonin. J Neurophysiol, 2008. 99(5): p. 2077–85.

90. Dacks, A.M., et al., Serotonin modulates olfactory processing in the antennal lobe of Drosophila. J Neurogenet, 2009. 23(4): p. 366–77.

91. Hardy, A., et al., 5-Hydroxytryptamine action in the rat olfactory bulb: in vitro electrophysiological patch-clamp recordings of juxtaglomerular and mitral cells. Neuroscience, 2005. 131(3): p. 717–31.

92. Kloppenburg, P., B.S. Kirchhof, and A.R. Mercer, Voltage-activated currents from adult honeybee (Apis mellifera) antennal motor neurons recorded in vitro and in situ. J Neurophysiol, 1999. 81(1): p. 39–48.

93. Mercer, A.R., J.H. Hayashi, and J.G. Hildebrand, Modulatory effects of 5-hydroxytryptamine on voltage-activated currents in cultured antennal lobe neurones of the sphinx moth Manduca sexta. J Exp Biol, 1995. 198(Pt 3): p. 613–27.

94. Mercer, A.R., B.S. Kirchhof, and J.G. Hildebrand, Enhancement by serotonin of the growth in vitro of antennal lobe neurons of the sphinx moth Manduca sexta. J Neurobiol, 1996. 29(1): p. 49–64.

95. Appel, N.M., et al., Autoradiographic characterization of (+-)-1-(2,5-dimethoxy-4-[125I] iodophenyl)-2-aminopropane ([125I]DOI) binding to 5-HT2 and 5-HT1c receptors in rat brain. J Pharmacol Exp Ther, 1990. 255(2): p. 843–57.

96. Hellendall, R.P., et al., Prenatal expression of 5-HT1C and 5-HT2 receptors in the rat central nervous system. Exp Neurol, 1993. 120(2): p. 186–201.

97. McLean, J.H., A. Darby-King, and G.D. Paterno, Localization of 5-HT2A receptor mRNA by in situ hybridization in the olfactory bulb of the postnatal rat. J Comp Neurol, 1995. 353(3): p. 371–8.

98. Shen, Y., et al., Molecular cloning and expression of a 5-hydroxytryptamine7 serotonin receptor subtype. J Biol Chem, 1993. 268(24): p. 18200–4.

99. Tecott, L.H., A.V. Maricq, and D. Julius, Nervous system distribution of the serotonin 5-HT3 receptor mRNA. Proc Natl Acad Sci U S A, 1993. 90(4): p. 1430–4.

100. Waeber, C., et al., Putative 5-ht5 receptors: localization in the mouse CNS and lack of effect in the inhibition of dural protein extravasation. Ann N Y Acad Sci, 1998. 861: p. 85–90.

101. Watts, S.W., et al., Autoradiographic comparison of [125I]LSD-labeled 5-HT2A receptor distribution in rat and guinea pig brain. Neurochem Int, 1994. 24(6): p. 565–74.

102. Brill, J., et al., Serotonin increases synaptic activity in olfactory bulb glomeruli. J Neurophysiol, 2015: p. jn 00847 2015.

103. Kapoor, V., et al., Activation of raphe nuclei triggers rapid and distinct effects on parallel olfactory bulb output channels. Nat Neurosci, 2016.

104. Liu, S., et al., Serotonin modulates the population activity profile of olfactory bulb external tufted cells. J Neurophysiol, 2012. 107(1): p. 473–83.

105. Huang, Z., N. Thiebaud, and D.A. Fadool, Differential serotonergic modulation across the main and accessory olfactory bulbs. J Physiol, 2017. 595(11): p. 3515–3533.

106. Lottem, E., M.L. Lorincz, and Z.F. Mainen, Optogenetic Activation of Dorsal Raphe Serotonin Neurons Rapidly Inhibits Spontaneous But Not Odor-Evoked Activity in Olfactory Cortex. J Neurosci, 2016. 36(1): p. 7–18.

107. Kaas, J.H., Topographic maps are fundamental to sensory processing. Brain Res Bull, 1997. 44(2): p. 107–12.

108. Kaneko, T. and B. Ye, Fine-scale topography in sensory systems: insights from Drosophila and vertebrates. J Comp Physiol A Neuroethol Sens Neural Behav Physiol, 2015. 201(9): p. 911–20.

109. Patel, G.H., D.M. Kaplan, and L.H. Snyder, Topographic organization in the brain: searching for general principles. Trends Cogn Sci, 2014. 18(7): p. 351–63.

110. Rothschild, G. and A. Mizrahi, Global order and local disorder in brain maps. Annu Rev Neurosci, 2015. 38: p. 247–68.

111. Gomez, C., et al., Heterogeneous targeting of centrifugal inputs to the glomerular layer of the main olfactory bulb. J Chem Neuroanat, 2005. 29(4): p. 238–54.

112. Gracia-Llanes, F.J., et al., Synaptic connectivity of serotonergic axons in the olfactory glomeruli of the rat olfactory bulb. Neuroscience, 2010. 169(2): p. 770–80.

113. Muzerelle, A., et al., Conditional anterograde tracing reveals distinct targeting of individual serotonin cell groups (B5-B9) to the forebrain and brainstem. Brain Struct Funct, 2016. 221(1): p. 535–61.

114. Lizbinski, K.M., et al., The anatomical basis for modulatory convergence in the antennal lobe of Manduca sexta. J Comp Neurol, 2016. 524(9): p. 1859–75.

115. Shang, Y., et al., Excitatory local circuits and their implications for olfactory processing in the fly antennal lobe. Cell, 2007. 128(3): p. 601–12.

116. Katz, P.S. and W.N. Frost, Intrinsic neuromodulation: altering neuronal circuits from within. Trends Neurosci, 1996. 19(2): p. 54–61.

117. Lizbinski, K.M. and A.M. Dacks, Intrinsic and Extrinsic Neuromodulation of Olfactory Processing. Front Cell Neurosci, 2018. 11: p. 424.

118. Dweck, H.K., et al., Olfactory preference for egg laying on citrus substrates in Drosophila. Curr Biol, 2013. 23(24): p. 2472–80.

119. Prieto-Godino, L.L., et al., Evolution of Acid-Sensing Olfactory Circuits in Drosophilids. Neuron, 2017. 93(3): p. 661–676 e6.

120. Semmelhack, J.L. and J.W. Wang, Select Drosophila glomeruli mediate innate olfactory attraction and aversion. Nature, 2009. 459(7244): p. 218-23.

121. Knaden, M., et al., Spatial representation of odorant valence in an insect brain. Cell Rep, 2012. 1(4): p. 392–9.

122. Fuyama, Y., Behavior genetics of olfactory responses in Drosophila. I. Olfactometry and strain differences in Drosophila melanogaster. Behav Genet, 1976. 6(4): p. 407–20.

123. Huoviala, P., et al., Neural circuit basis of aversive odour processing in Drosophila from sensory input to descending output. BioRxiv, 2019.

124. Silbering, A.F., et al., Complementary function and integrated wiring of the evolutionarily distinct Drosophila olfactory subsystems. J Neurosci, 2011. 31(38): p. 13357–75.

125. Cohen, J.Y., M.W. Amoroso, and N. Uchida, Serotonergic neurons signal reward and punishment on multiple timescales. Elife, 2015. 4.

126. Jacobs, B.L., J. Heym, and M.E. Trulson, Behavioral and physiological correlates of brain serotoninergic unit activity. J Physiol (Paris), 1981. 77(2-3): p. 431–6.

127. Monti, J.M., Serotonin control of sleep-wake behavior. Sleep Med Rev, 2011. 15(4): p. 269–81.

128. Seo, C., et al., Intense threat switches dorsal raphe serotonin neurons to a paradoxical operational mode. Science, 2019. 363(6426): p. 538-542.

129. Bunin, M.A. and R.M. Wightman, Quantitative evaluation of 5-hydroxytryptamine (serotonin) neuronal release and uptake: an investigation of extrasynaptic transmission. J Neurosci, 1998. 18(13): p. 4854–60.

130. Bunin, M.A. and R.M. Wightman, Paracrine neurotransmission in the CNS: involvement of 5-HT. Trends Neurosci, 1999. 22(9): p. 377–82.

131. Fu, W., et al., Chemical neuroanatomy of the dorsal raphe nucleus and adjacent structures of the mouse brain. J Comp Neurol, 2010. 518(17): p. 3464–94.

132. Liu, Z., et al., Dorsal raphe neurons signal reward through 5-HT and glutamate. Neuron, 2014. 81(6): p. 1360–74.

133. Sengupta, A., et al., Control of Amygdala Circuits by 5-HT Neurons via 5-HT and Glutamate Cotransmission. J Neurosci, 2017. 37(7): p. 1785–1796.

134. Gasque, G., et al., Small molecule drug screening in Drosophila identifies the 5HT2A receptor as a feeding modulation target. Sci Rep, 2013. 3: p. srep02120.

135. Nichols, D.E. and C.D. Nichols, Serotonin receptors. Chem Rev, 2008. 108(5): p. 1614–41.

136. Zhang, B., M.R. Freeman, and S. Waddell, Drosophila neurobiology: a laboratory manual. 2010, Cold Spring Harbor, N.Y.: Cold Spring Harbor Laboratory Press. x, 534 p.

137. Saalfeld, S., et al., CATMAID: collaborative annotation toolkit for massive amounts of image data. Bioinformatics, 2009. 25(15): p. 1984–6.

138. Schneider-Mizell, C.M., et al., Quantitative neuroanatomy for connectomics in Drosophila. Elife, 2016. 5.

139. Li, P.H., et al., Automated Reconstruction of a Serial-Section EM Drosophila Brain with Flood-Filling Networks and Local Realignment. BioRxiv, 2019.

140. Milyaev, N., et al., The Virtual Fly Brain browser and query interface. Bioinformatics, 2012. 28(3): p. 411–5.

141. Bates, A.S., et al., The natverse: a versatile computational toolbox to combine and analyse neuroanatomical data. BioRxiv, 2019.

142. MacQueen, B., Some methods for classification and analysis of multivariate observations. Proceedings of 5-th Berkeley Symposium on Mathematical Statistics and Probability, 1967. 1: p. 281-297.

143. Rousseeuw, P., Silhouettes: a Graphical Aid to the Interpretation and Validation of Cluster Analysis. Computational and Applied Mathematics., 1987. 20: p. 53–65.

